# Simultaneous cortical responses to multiple written words

**DOI:** 10.1101/2025.06.05.658099

**Authors:** Vassiki Chauhan, Krystal McCook, Mariam Latif, Alex White

## Abstract

We have learned much about the brain regions that support reading by measuring neuronal responses to single words, but we know little about how the brain processes multiple words simultaneously. This fMRI study fills that gap by varying the number of English words presented while holding the amount of visual stimulation constant. We adapted the “simultaneous suppression” paradigm, which has demonstrated that the response to multiple stimuli presented simultaneously is typically smaller than the sum of responses to the same stimuli presented sequentially. On each trial, participants viewed a rapid sequence of three frames. Each frame contained two character strings, most of which were pseudo-letters with visual features matched to familiar letters. The experimental conditions differed in the number of words in the sequence: zero words; one word; two words sequentially; or two words simultaneously. Behavioral task accuracy was worse for detecting two words presented simultaneously than sequentially. BOLD responses increased linearly with the number of words presented in several reading-related regions of the left hemisphere: text-selective occipito-temporal regions, the superior temporal sulcus, the intraparietal sulcus, and the inferior frontal sulcus. In all of those regions, responses did not differ between sequential and simultaneous presentation of two words. Nonetheless, the sensitivity of ventral temporal text-selective regions to the lexical frequencies of two words was attenuated by simultaneous presentation. To account for these patterns of activity and task performance, we suggest that the reading network can detect two strings of letters simultaneously, but there is interference during lexical access.

## INTRODUCTION

To read this page, you must integrate information across many hierarchically arranged visual elements, from letters to words to sentences. Fluent reading requires a balance between holistic parallel processing of multiple elements and serial processing of chunks of text to avoid interference. Here, we investigate how reading-related brain areas process multiple words simultaneously.

Most fMRI studies of visual word recognition have presented only one word at a time (e.g., Caffarra et al., 2021; Dehaene & Cohen, 2011; Yeatman & White, 2021). Others analyzed activity during sentence reading (e.g., Carter et al., 2019; Henderson et al., 2015; Schuster et al., 2015) but were designed for other questions. EEG and MEG studies have provided important insights into the dynamics of multi-word processing (e.g., Dunagan et al., 2025; Fallon & Pylkkänen, 2024; Flick et al., 2021; Schotter et al., 2023; Wang et al., 2025), but fMRI offers superior spatial resolution for specific reading-related regions.

Behavioral studies have raised deep questions about how efficiently readers can process multiple words in parallel. The *parafoveal preview benefit*, for example, demonstrates that while readers fixate on word N, they begin processing word N+1 (Schotter et al., 2012; Vasilev & Angele, 2017). Some models of reading account for such effects by allowing parallel processing of multiple words with divided attention (Engbert et al., 2005; Reilly & Radach, 2006; Snell et al., 2018; Snell & Grainger, 2019). Other models assume that readers finish processing the fixated word before shifting attention covertly to the next word(s) (Pollatsek et al., 2006; Schotter et al., 2014). Several experimental tasks have also provided support for serial models (White et al., 2018, 2020; Brothers et al., 2022; Campbell et al., 2024; Johnson et al., 2022).

Behavioral deficits might arise from limited processing capacity in the “visual word form area” (VWFA), a left ventral occipito-temporal region that links the visual system and the language system (Dehaene & Cohen 2011; Yeatman & White, 2021). White et al. (2019) argued that a posterior VWFA sub-region can process two words at once in separable “channels”. In contrast, activity in the anterior sub-region was affected by the lexical frequency of only one of two words per trial, consistent with a processing ‘bottleneck’ during lexical access (see also White et al. 2020). However, that study did not include a comparison condition with words presented one at a time.

Here we leverage another neural correlate of processing capacity limits in vision: *simultaneous suppression*: a lower BOLD response to a set of stimuli flashed simultaneously compared to the response to the same stimuli flashed in rapid sequence (Kastner et al., 1998; Kastner & Ungerleider, 2001; McMains & Kastner, 2011). This phenomenon supports theories that multiple stimuli “compete” for processing resources (Miller et al., 1993; Desimone & Duncan, 1995; Beck & Kastner, 2009). Ours is the first study to draw on this literature to investigate how the responses in text- and language-selective regions vary as a function of the number of words being read.

Previously reported simultaneous suppression effects could be due to more stimulus onset and offsets in the sequential condition, each of which causes a transient neural response, rather than the competitive interactions between stimuli (Kupers et al., 2024). Therefore, we kept visual stimulation constant across all conditions by inserting zero, one, or two words into rapid streams of “false font” strings with similar visual features. The first question is whether BOLD responses (relative to baseline with no stimulation) increase steadily as the number of words increases. The second question is whether the response to two words presented simultaneously is as strong as the summed response to two words presented sequentially. We also compare behavioral performance across those conditions, similar to previous studies that have argued for limited processing capacity (e.g., Scharff et al 2011; White et al. 2026). Following White et al. (2019), we also analyze activity as a function of the lexical frequencies of multiple words.

## METHODS

### Participants

We recruited 22 participants (5 self-identified as male) from the Barnard College and Columbia University student body, and general New York City Community (N = 4). As described below, one participant had to be excluded from the analysis due to excessive head motion, leaving a final sample of 21 participants. Their ages ranged from 19 to 40 years (mean = 24.4土6.3 years). All but one were right-handed. All participants had learned English before the age of five, and provided written informed consent. They were monetarily compensated for their participation. The study was approved by the Internal Review Board at Barnard College.

The sample size was chosen in advance of data collection. In our two previous fMRI studies of BOLD responses to written words with similar protocols (Chauhan et al., 2024; White et al., 2023), we included 15 and 17 participants, respectively. In this study, we explore a novel effect (simultaneous suppression for words) which we expected might be smaller than the task effects detected in those prior studies. We therefore set out to collect as much data as we had funding for, which allowed full data sets for 22 participants.

The sample size is also justified by a power analysis of the theoretical simultaneous suppression effect in the VWFA, using data from Chauhan et al. (2024). In that study, we estimated the left VWFA’s response magnitude to single words and false font strings. We assumed that in this new study, the response to single words would have the same mean (0.32 percent signal change) and standard deviation (0.2) as in the previous study. (The new data matched that prediction quite closely: mean = 0.30 p.s.c, SD = 0.18). We then assumed that the response in the two-word sequential condition would be equal to the 1-word mean plus the difference between the response to single words and false font strings in Chauhan et al. (2024). This is because both conditions include responses to false font strings. According to a serial processing model, the response to the two-word *simultaneous* condition would be the same as in the one-word condition. That would be the case if a “serial bottleneck” allows only one of the two simultaneous words to be processed, as some of our previous work concluded (White et al., 2019). We then simulated responses in the 2-simultaneous and 2-sequential conditions, using those predicted means and standard deviations. To reject the null hypothesis for the difference between those two conditions (i.e., the simultaneous suppression effect) with 90% power, 18 participants would be sufficient. Rather than stop after that number of participants, we planned in advance to collect as much data as we could, which left us with 22 participants (one of which had to be excluded for excessive head motion).

### Equipment

We acquired MRI data at the Zuckerman Institute, Columbia University. We used a 3T Siemens Prisma scanner and a 64-channel head coil. At the start of each scanning session, we acquired a T1-weighted structural scan, with 1 mm isometric voxels. We collected functional data with T2* echo planar imaging sequences with multiband echo sequencing (SMS3) for whole brain coverage without compromising temporal resolution. The TR was 1.5 s, TE was 30 ms, the flip angle was 62° and the voxel size was 2 mm isotropic.

We presented the stimuli using custom code written in MATLAB and the Psychophysics Toolbox (Brainard, 1997). We projected stimuli onto a screen that the participants viewed through a mirror, with a viewing distance of 142 cm. The resolution of the display was 1920 by 1080 pixels, and the refresh rate was 60 Hz. We also collected eye-tracking data with an SR Research Eyelink 1000 tracker. To respond to the tasks, the participant used their right hand to press one of three buttons on a handheld MR-safe response pad.

### Stimuli

The stimuli and task for the main experiment are depicted in **Figure 1A**. A small black fixation dot was continuously present at the center of the screen against a light background (95% of screen maximum). The dots diameter was 0.11 degrees of visual angle (dva). The stimuli were five-letter English words. We selected 768 such words with no constraints on parts of speech. On the Zipf scale, the lexical frequencies ranged from 2.2 to 6.1, with a median of 4.1. Zipf is a standardized measure of word frequency calculated through an adjustment of the log frequency per million words in the SUBTLEX-US corpus (Brysbaert & New, 2009; van Heuven et al., 2014).

**Figure 1:**
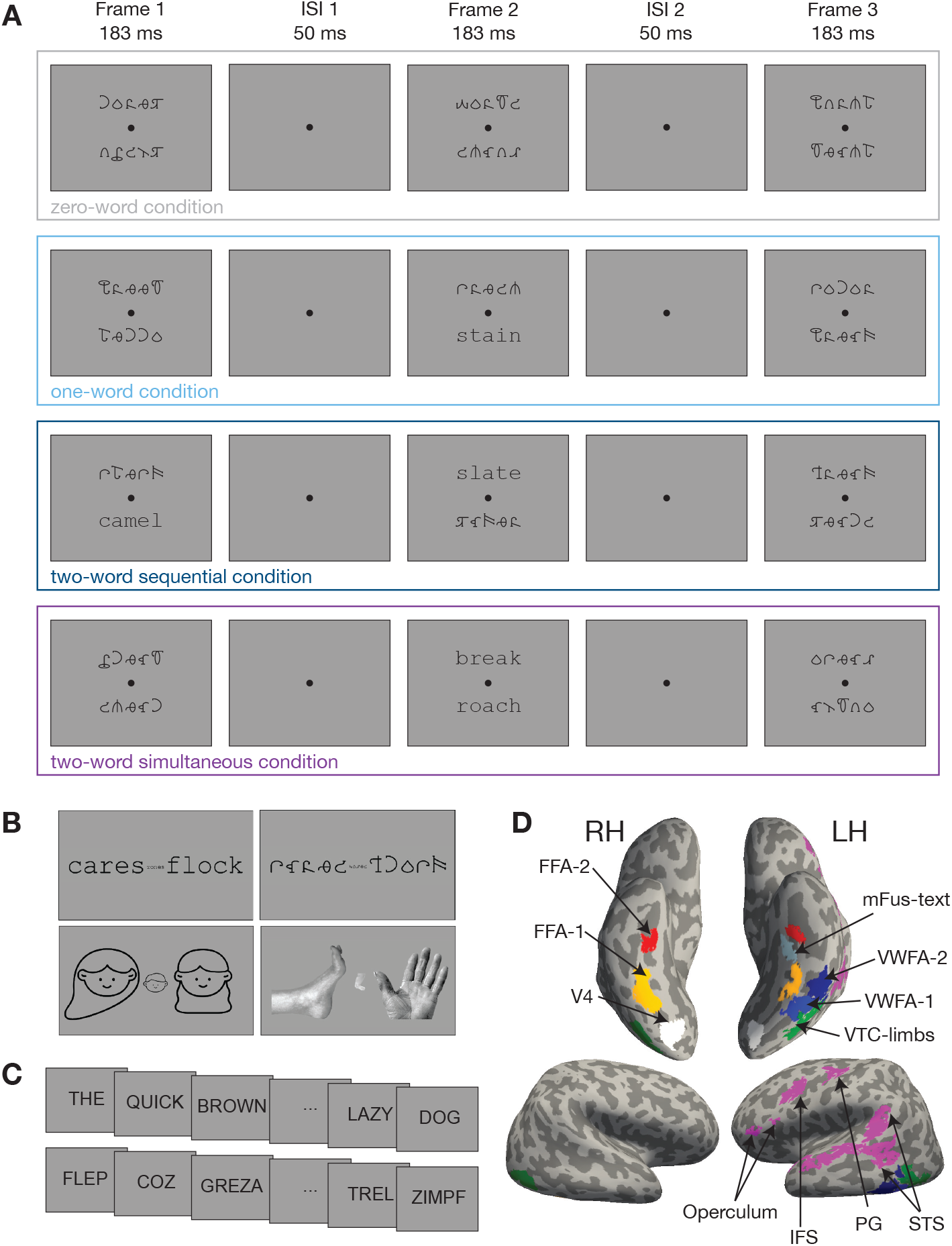
Example stimuli and regions of interest (ROIs). (**A**) Example trials of each presentation condition in the main experiment. The four conditions differed in whether words appeared in the first and/or second frame. Frame 3 always contained two false font strings. The timing and location of words was counterbalanced within each condition (e.g., on one-word trials the word could be in either frame 1 or 2, on the top or bottom). After frame 3, the participant had 3.65 s to make their response. **(B)** Examples of the four stimulus categories used in the visual category-selective region localizer (words, false font strings, faces, and limbs). **(C)** Example stimuli from the language localizer. Each real sentence consisted of 12 real English words, and each jabberwocky sentence consisted of 12 pseudowords. Each (pseudo)word was presented for 450 ms followed by a 50 ms blank. **(D)** Example ROIs in a representative subject’s inflated cortical surfaces. The regions on the ventral view (top two images) were defined from the visual category localizer. The white V4 region was obtained from a previous study (White et al., 2023). The pink regions on the lateral view (bottom images) were defined from the language localizer. Abbreviations: FFA: fusiform face area; pOTS-words: word-selective posterior occipitotemporal sulcus; mOTS-words: word-selective middle occipitotemporal sulcus; mFus-words: word-selective mid-fusiform sulcus/gyrus; VTC-limbs: limb area in the ventral temporal surface, extending laterally to include the extrastriate body area (EBA) in some subjects; STS: superior temporal sulcus; PG: Precentral Gyrus; IFS: inferior frontal sulcus.

Real words were presented in the Courier New, a standard monospaced serif font. The other “filler” strings were presented in an unfamiliar, illegible false font called BACS-2 (Vidal et al., 2017). These false font characters are designed to match the size, the symmetry, and the numbers of strokes, junctions and terminations of Courier New. Both these fonts were scaled so that the height of the letter ‘x’ was 0.41 dva. On average, the width of each character string was 2.65 dva, and the height was between 0.41 and 0.79 dva. The distance between centers of neighboring letters was 0.55 dva. The stimuli were rendered in the same black color as the fixation dot.

### Trial sequence and task

The stimulus presentation on each trial consisted of three successive frames. Two character strings were presented in each frame, one above and one below the fixation dot. The centers of the character strings were 1.5 dva from the fixation dot. This display arrangement differs from the conditions of natural English reading, but we chose it for two reasons: because behavioral evidence for serial processing of words is strong with that arrangement (White et al., 2020), and because both words are equally near to fixation and clearly legible. Arranging the words horizontally would better match the conditions of natural reading, but there is a strong right>left hemifield asymmetry (Yeatman & White, 2021). Indeed, a strong bias to process words on the right complicated the results of White et al (2019). Crucially, the VWFA is known to respond strongly to words in a region spanning roughly the central 5 degrees of the visual field (R. K. Le et al., 2017; White et al., 2023), including just above and below fixation (Rauschecker et al., 2012). Thus, the population receptive fields (PRFs) of the voxels we analyze are known to cover both of the stimulus positions.

Each frame lasted for 183 ms, and there was a 50 ms inter-stimulus interval between frames (with only the fixation dot presented).

Across the three frames in each trial, there were always 6 character strings, and all of those that were not legible words were in fact English words drawn from the same set, but presented in the illegible false font. The number of words presented in each trial varied across four conditions, which were randomly intermixed. Examples of each condition are illustrated in **Figure 1A**. They are:

1. Zero-word trials: all the character strings were false fonts.
2. One-word trials: one real English word in Courier New was presented in either the first or second frame, either above or below fixation (equally likely).
3. Two-word sequential trials: one word was presented at one location in the first frame, and a different word was presented in the other location in the second frame. The first word was equally likely to be at the top or bottom location.
4. Two-word simultaneous trials: two words appeared at the same time, one above and one below the fixation, in either the first or the second frame (equally likely).

The third frame always contained two false font strings and served as a post-mask. After the third frame, there was a 3651 ms inter-trial interval in which the participant could press a button with their right hand to make their response. Their task was to report the number of words that had appeared in that trial (0, 1 or 2). The distinction between sequential and simultaneous presentation of two words was not relevant to the task. Participants used their ring finger to report zero words, middle finger for one word and index finger for two words. They were instructed to prioritize accuracy over speed. Participants were also instructed to read the words silently in their head.

After every block of five trials, there was a blank period, the duration of which was randomly and uniformly selected to be 4, 6 or 8 s. There were 64 trials in each run, presented in random order.

Each legible word was presented exactly once to each participant during the experiment. Orthographic neighborhood size and log lexical frequency estimated from the English Lexicon Project were balanced across all experimental conditions in order to minimize differences in task difficulty (Balota et al., 2007). We created 6 sets of word lists that balanced these statistics across the four conditions. One list was randomly chosen for each participant. Overall, 4 unique participants saw lists 1-4, and 3 unique participants saw lists 5 and 6.

The main experiment consisted of 8 five-minute fMRI runs. In total, there were 128 trials of each of the four main conditions. Each run began with written instructions, reminding the participants which fingers to use to make their responses, to respond as accurately as possible, to maintain central gaze fixation, and to read each word silently aloud in their head. There was no feedback at the end of each trial, but the percent correct was displayed once the run ended, along with the percentage of trials in which the participants failed to make a response on time.

Prior to scanning, participants practiced the task in a testing room outside the scanner. During practice, they used the same fingers to report the number of words as during scanning, and their gaze position was tracked. On these practice trials they received auditory feedback: a high-pitched tone for the correct response and a low-pitched tone for the incorrect response. They performed at least 32 practice trials, and repeated the block if their accuracy was below 80%. They also viewed the stimuli for the localizer experiments during the practice session.

### Functional Localizers

Each participant completed two functional localizers: one to define category-selective ventral temporal visual regions, and one to localize core language regions. The visual category localizer is described in White et al. (2023) and Chauhan et al. (2024) (see also https://github.com/Barnard-Vision-Lab/catLoc). Briefly, participants viewed images of four categories of objects: text (including English words and pseudowords), false font strings, faces, and limbs. See examples in **Figure 1B**. These categories were chosen to maximize contrast between activity in the VWFA and surrounding face- and body-selective regions. Each frame included three examples of each category, one at fixation and two larger ones, one to either side. We used two fonts for the text, Courier New and Sloan, and two false fonts with matched visual features, BACS-2 (Vidal et al., 2017) and PseudoSloan (Vildavski et al., 2021). Each 4-second trial consisted of four frames with images of all the same category. Participants performed a one-back repetition detection task, pushing a button with their index finger whenever they saw the same exact image repeated two times in a row. Repetitions occurred on 33% of trials. Each participant completed four runs, each of which consisted of 54 trials and lasted for about six minutes. Each category was presented 56 times.

The second localizer was the language localizer scan, example trials of which are shown in **Figure 1C**. It was designed to define language processing centers in the frontal lobe and superior temporal lobe. See Fedorenko et al. (2010) for details. Briefly, the stimuli consisted of written words presented one at a time, 450 ms per word, 50 ms interstimulus interval between individual words. In the sentence condition, a sequence of 12 real English words formed a meaningful and grammatical sentence. In the “nonsense” condition, a sequence of 12 pronounceable pseudowords formed a meaningless “Jabberwocky” sentence. After every set of 12 character strings, an icon of a hand appeared. Participants’ task was simply to press a button with their index finger when they saw that hand, to ensure that they were paying attention to the stimuli. Participants performed two runs of this task, each of which consisted of sixteen 6-s trials. Each run also included five blank trials which also lasted 6 s.

Regions of Interest defined using both localizers are depicted in **Figure 1D**.

### Study Procedure

For 18 out of the 22 subjects, the entire study required three sessions. The first session involved training the participants to perform the tasks and calibrate the eye-tracker. The other two sessions were conducted in the MRI scanner and both included the same sequence of types of scans. First, while we acquired a T1 structural scan, participants performed a few practice trials. Then they performed 4 runs of the main experiment, 2 runs of the visual category localizer and 1 run of the language localizer. The other 4 participants had already completed the localizer scans in a previous study, so they only performed 8 runs of the main experiment during a single MRI session.

### Analysis of Behavior

We defined accuracy as the proportion of trials with correct responses made within the response period, which lasted 3.65 s after the offset of the final stimulus frame. On average, participants failed to respond on time on 1.3% of trials (SEM=0.6%). Those trials were excluded from the analysis of accuracy.

### Eye-tracking data analysis

We used an Eyelink 1000 eye-tracker (SR Research) to record gaze position data during each scan. However, because of a suboptimal placement of the camera and infrared illuminator, the data quality varied across participants. In some cases, it was not possible to calibrate the tracker, or the tracker periodically failed to locate the participant’s pupil. For two participants, we could not collect any gaze data.

The first step in our analysis of the eye-tracking data was to determine which trials had data of acceptable quality. A whole run was excluded from the analysis of gaze positions if the median standard deviation of gaze positions 250 ms before stimulus presentation was greater than 1°. We chose that interval because participants were usually fixating stably just before the stimuli arrived. Rapidly changing position estimates during that interval are a sign of a poor data quality. 5 runs were excluded from the eye-tracking analysis for this reason (from a total of 3 participants).

Next, within included runs, each trial was deemed analyzable if, during the 650 ms of stimulus presentation, at least 80% of the gaze position samples were acceptable. Unacceptable samples are those with missing position values or positions >10° from the fixation mark, which is implausible and a sign of calibration failure. After applying those criteria, we had acceptable gaze data on 51% of trials. For 8 of the participants, more than 95% of trials were acceptable, and the rest covered a wide range.

For the trials with acceptable gaze data, we detected fixation breaks as follows: We first conducted a drift correction within each run. Specifically, across all trials in the run, we computed the median horizontal and vertical gaze position in the 250 ms before stimulus onset. Then, we defined the corrected fixation position as the across-trial median of those medians, including only trials in which at least 90% of the eye position samples in the pre-stimulus interval were acceptable.

Next, we analyzed the gaze position recorded during the 650 ms of stimulus presentation on each trial, calculating the distance of each gaze sample from the corrected fixation position. We defined a fixation break as a deviation of more than 1.5° from the corrected fixation position for at least 100 ms, or if the participant blinked (as evidenced by pupil size dropping to 0 for at least 100 ms during the stimuli). We also detected saccades using the method described by Engbert & Mergenthaler (2006). Saccades were events in which the two-dimensional gaze position velocity exceeded, for at least 6 ms, an ellipse with horizontal and vertical radii equal to 4 times the horizontal and vertical median-based standard deviations, respectively. Those standard deviations were calculated in the interval from 250 ms prior to stimulus onset until stimulus offset. Saccades had to be >1° and <15° in amplitude. We also categorized individual trials as fixation breaks if the vertical distance of the saccade exceeded 1.5°.

With those criteria, fixation breaks occurred on 4.95% of trials with acceptable gaze data (ranging across participants from 0 to 18.8%). Saccades with a vertical amplitude more than 1.5° occurred on 2.16% of trials (range 0 to 17.0%).

To go further, we also estimated how often a word was fixated on directly for at least 50 ms (that is, the gaze position was within the word’s bounding box during the 183 ms it appeared). This occurred on only 0.17% of trials with acceptable gaze data and at least 1 word present (and never for 17 of the participants).

Thus, within the runs with acceptable gaze data, we conclude that participants were fixating well. Fixation breaks occurred on less than 5% of trials overall. Saccades were rare, and individual words hardly ever got fixated directly. We also found little evidence that the appearance of a word attracted the gaze to its side, even in the next frame.

### Preprocessing

We pre-processed our MRI data with *fMRIPrep* 21.0.1 (Esteban et al., 2019), which is based on *Nipype* 1.6.1 (Gorgolewski et al., 2011). Depending on the number of sessions, we obtained one or two T1-weighted structural scans for each individual. These structural images were skull-stripped, corrected for intensity non-uniformity, and averaged across sessions for participants who participated in multiple sessions. Cortical surfaces were constructed using the boundaries between gray matter and white matter using Freesurfer (Fischl, 2012).

The functional MRI scans were preprocessed as follows: Magnetic field inhomogeneities in each EPI sequence were corrected using a B_0_ nonuniformity map collected during the same session. All functional runs were co-registered to the native anatomical volume (*fsnative*) We then slice-time corrected the multiband sequences and resampled them onto each participant’s cortical surface.

We also used fMRIPrep to perform motion correction. Moreover, in order to ensure that our data would not be contaminated by head motion, we used the following exclusion criteria. A run with movement greater than 2 voxels (4 mm) was excluded from analysis. Additionally, if any participant had such excessive head motion in more than half of the runs, that participant was dropped from the analysis. Using these criteria, we excluded one participant from the analysis entirely, leaving a final N of 21. A total of three runs were excluded for head motion from 2 of the included participants.

### BOLD Response Estimation

We estimated BOLD responses (beta weights) to each single trial with GLMSingle in Python (Prince et al., 2022). The result of this analysis is a beta weight for each trial, for each cortical surface node in each participant. We first resampled the BOLD time-series from 1.5s to 1s. GLMSingle optimizes the hemodynamic response function for each node in the surface space and performs denoising using fractional ridge regression to remove correlated noise. For the main experiment, the design matrix consisted of a unique column for each stimulus condition (zero, one, two-sequential or two-simultaneous words), uniquely coding trials in which the top word appeared in the first or second frame, and uniquely coding the frame in which the words in the 1-word or 2-simultaneous conditions appeared. For the visual category localizer, each stimulus category received its own column. For the language localizer, there were two columns: sentence trials and nonsense trials.

### ROI Definition

We sought to define nine regions of interest (ROIs) in each hemisphere in each participant. See **Figure 1D** for these ROIs on an example participant’s inflated cortical surfaces. Details about the number of participants in which we were able to localize each ROI are provided in the tables that report statistics on the stimulus effects in each ROI. Six of the ROIs in the ventral temporal cortex were obtained from the visual category localizer scans. For each category (words, faces and limbs), we computed t-statistics for the difference in BOLD responses to that category vs. all other categories (including false fonts). The t-values were projected onto each participant’s inflated cortical surfaces, and thresholded at t ≥ 3.

We defined up to three text-selective areas in each hemisphere using the following anatomical and functional constraints. We defined two text-selective areas in or on the borders of the occipitotemporal sulcus: *pOTS-words* was in the posterior part of the sulcus, but anterior to the visual area hV4; *mOTS-words* was in the middle of the sulcus. In some subjects, there was a continuous patch that we segmented into mOTS-words and pOTS-words. These regions have also been labeled “VWFA-1” and “VWFA-2” (e.g., White et al., 2019; Chauhan et al., 2024; Yablonski et al., 2024). Here we prefer to use the anatomical labels, in line with other authors (Dalski et al., 2024; Grill-Spector & Weiner, 2014; Lerma-Usabiaga et al., 2018). A third text-selective region was *mFus-words*, in the middle of the fusiform gyrus or mid-fusiform sulcus, medial to and often anterior of mOTS-words. The face-selective areas, FFA-1 and FFA-2, were always medial to the text-selective OTS regions and located on the fusiform gyrus. The limb-selective area was lateral pOTS and mOTS-words, and in some subjects extended into posterior inferior temporal sulcus/middle temporal gyrus (the location of the putative “extrastriate body area”, Downing et al., 2001)

We also used the language localizer to define four core language regions with the contrast of sentences vs. nonsense sequences. These language ROIs were in the precentral gyrus (PG), the inferior frontal sulcus (IFS), the frontal operculum, and the superior temporal sulcus (STS).

Lastly, we obtained one bilateral retinotopic visual region, hV4, from a previous study that used retinotopic mapping (White et al., 2023). This average-subject ROI was projected from the *fsaverage* template into each participant’s native surface space.

### Statistical Analysis of Behavior and ROI Responses

We used linear mixed effects modelling to analyze both behavioral and brain responses on single trials. To analyze the effect of presentation condition on task accuracy, a generalized linear model with a binomial linking function predicted the probability of a correct report of the number of words, *pCorr*, as a function of the presentation condition *C*:

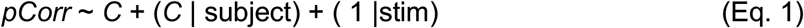

The presentation condition *C* (0, 1, 2-sequential or 2-simultaneous words) was a fixed effect. We also included random slopes and intercepts by subject, and random effects for the stimuli (“stim”) presented on each trial. To code the stimuli on each trial as a single variable we used the following procedure: for two-word trials, the stimulus was coded as the concatenation of the two words. For one-word trials, the stimulus was coded as that one word. For zero-word trials, which only included false fonts, the item was coded as “{string}_ff”, where {string} was a randomly selected one of the four false font strings presented in the first two frames on that trial. This string was in fact a word but presented in the illegible false font. We assume that our participants (nor any of the brain regions of interest) cannot reliably distinguish these false font strings from each other, so the choice of items to code the 0-word trials variable was arbitrary. Thus, all trials had a unique character string that coded the stimuli on that trial, which was necessary for the models to converge.

The calculation of correct response times (RTs) was somewhat complex, given that the stimuli were spread out over multiple frames that were offset in time by 233 ms. Therefore, for each trial, we calculated response time starting from the onset of the “critical frame,” which was the frame that disambiguated the correct choice. Frame 2 was the critical frame for all trials except those with two words simultaneously in frame 1. To ensure an unbiased comparison between conditions, we excluded from analyses of RTs two-word trials with both words in frame 1, and 1-word trial with the word in frame 1. Thus, all the RTs we analyzed were calculated from the onset of frame 2, and all (except the 0-word condition) had at least one word appearing in frame 2. However, the qualitative pattern of RT differences across conditions was the same even if we did not exclude any trials. RTs were fit by the following linear mixed effects model:

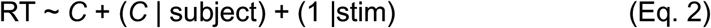

To analyze BOLD responses within each ROI, we used two different linear mixed effects models to predict the beta weight (*β)* on each trial. We used the *fitlme* function in MATLAB for these analyses. The first model tested the ***number of words effect***: how much the BOLD response increased linearly from 0 to 1 to 2-word trials:

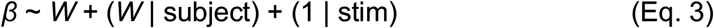

The number of words, *W*, (0, 1 or 2) was a numeric fixed effect, and the model included random slopes and intercepts across subjects and a random effect for stimulus items (“stim”, as explained above). We excluded the two-word simultaneous condition because a goal of this first analysis was to verify that the BOLD magnitude represented an integration of the response to two sequential words. We then ran two pairwise contrasts: 0 vs 1 word, and 1 vs 2 words.

The second model tested the ***simultaneous suppression effect***, the comparison of sequential vs. simultaneous presentation conditions only for the two-word trials. This model was similar to the previous except that the fixed effect was the presentation condition, *C*, as a categorical variable.

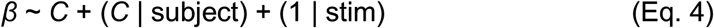

To obtain p-values, the outputs from the linear mixed effects model calls needed to be transformed to the ANOVA format, which we report along with the accompanying F-statistic in the manuscript. The degrees of freedom (DF) were calculated as the difference between the number of observations in the single trial data and the number of fixed effects. For each test, we applied a false discovery rate (FDR) correction to the p-values across regions of interest for a false discovery rate of 0.05 (Benjamini & Hochberg, 1995).

To calculate Bayes Factors for each effect and interactions between conditions, we used the MATLAB bayesFactor toolbox (Krekelberg, 2024). This toolbox required us to first average the responses over trials for each participant, and did not allow random slopes by subject. Each Bayes Factor (BF_10_) is the ratio of the probability of the data under the alternate hypothesis (a distance is >0 or two conditions differ) relative to the probability of the data under the null hypothesis that there is no difference (Rouder et al., 2009). A BF of 10 indicates that the data are 10 times more likely under the alternate hypothesis than the null. A BF of 0.10 indicates the opposite.

Our primary ROIs are the three text-selective sub-regions on the ventral temporal cortex (identified with the words vs. other categories contrast). The VWFA contains multiple sub-regions that might have different functions. We refer to two patches within the left occipito-temporal sulcus: a posterior one called pOTS-words or VWFA-1, and a relatively anterior one called mOTS-words or VWFA-2. These two areas have different stimulus response patterns, functional connectivity, and white matter connectivity (Kubota et al., 2023; Lerma-Usabiaga et al., 2018; White et al., 2019; Yablonski et al., 2024; Yeatman & White, 2021). Intracranial recordings have revealed a reading-related site on the mid-fusiform gyrus that is anterior and medial of the “classical” VWFA (Woolnough et al., 2021). Within the ventral temporal cortex, this area is the first to respond differentially to low- vs high-frequency words. Woolnough and colleagues (2021) propose that the mid-fusiform area supports the brain’s “orthographic lexicon” (see also Taylor et al., 2013). In our study, we therefore localized a mid-fusiform region (“mFus-words”), in addition to pOTS-words and mOTS-words in the occipito-temporal sulcus.

For those regions, FDR correction was applied to each set of 3 p-values. We applied the same analyses to all the other visual and language ROIs described above, and across all of these we applied the FDR correction. The statistics in the text and the Tables report these corrected p-values. The pairwise contrasts are reported in the Results sections and the figures. We used reference dummy coding for the categorical fixed effects, and then applied planned contrasts using MATLAB’s *coefTest* function on the model generated by *fitlme* function to compare pairwise conditions (e.g., the mean BOLD response in the 1-word condition vs the 2-word sequential condition).

Moreover, to determine whether the patterns were consistent across the three text-selective ventral temporal areas, and to maximize power for detecting simultaneous suppression there, we also fit additional models that included “region” as a categorical fixed effect (pOTS-words, mOTS-words, or mFus-words) that interacted with the stimulus condition. For the number of words effect, the model was:

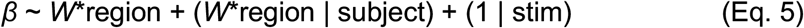

For the simultaneous suppression effect on two-word trials, the model was:

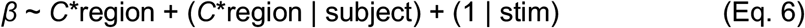

Finally, we examined the effect of the words’ lexical frequencies on behavior and brain responses in ventral text-selective areas on two-word trials. This analysis compared trials in which both words were low frequency (below median of the stimulus set) or both words were high frequency (above the median). To do so we fit similar mixed effect models as described above, with the addition of lexical frequency bin as a fixed effect (“*freqBin*”).

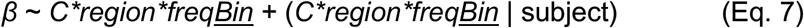

Individual stimuli could not be included as a random effect in this analysis because they were highly correlated with frequency bin. Behavioral effects of lexical frequency were modelled similarly as above:

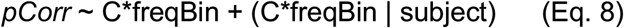

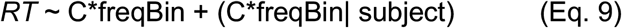

### Whole-cortex Analyses

We performed two whole-cortex analyses within each participant. The first was the “number of words effect”: a linear slope fit to beta weights as a function of the number of words presented in the 0, 1-word, and 2-words sequential conditions. The second was a contrast of the “simultaneous suppression” effect: the difference in beta weights between two-word sequential trials and two-word simultaneous trials. Therefore, for the number of words effect, each node was assigned a slope, and for the simultaneous suppression contrast, each node was assigned the difference in response to sequential and simultaneous trials. After computing each statistic in each subject’s native surface space, we spatially smoothed the data with a 5 mm full-width at half-maximum kernel, and projected the smoothed data onto *fsaverage* space using the *mri_surf2surf* function from Freesurfer. We then concatenated all participants’ contrasts in *fsaverage* space and computed the mean at each node. Next, we performed a one-sample t-test to determine the nodes in which the value was consistently different from zero. Finally, we corrected the associated p-values for multiple comparisons using FDR correction, implemented as *fdrcorrection* in the Python library *statsmodels.* We then saved the t-values for nodes with correct p<0.05.

## RESULTS

The experiment included four main conditions: zero-word trials, one-word trials, two-word sequential trials and two-word simultaneous trials (**Figure 1A**). The design allowed us to test two hypotheses: 1) That the response magnitude in reading-related regions increases with the number of items of their preferred stimulus category (i.e., words), while the amount of visual stimulation is held constant. 2) That the response to two simultaneously presented words is lower than the total response to two sequentially presented words, due to processing capacity limits. This second hypothesis is based on earlier studies of simultaneous suppression in visual cortex (e.g., Kastner, Weerd, et al., 2001; Kupers et al., 2024), and on prior behavioral studies that show limited capacity for processing two words at once (e.g. White et al. 2020, White et al. 2026). First, we report the analysis of task performance in the scanner.

### Behavioral performance shows a cost of simultaneous presentation of two words

The participant’s task during scanning was to report the number of words they perceived on each trial. The task was designed to motivate participants to maintain central fixation while attending covertly to both locations throughout each trial; words could appear unpredictably at either or both locations. The display was too fast to allow goal-directed saccades during a single frame (of 183 ms duration). A prior study provides behavioral evidence that participants can divide spatial attention in order to encode visual information from these locations, even if a higher-level bottleneck prevents *recognition* of both words simultaneously (White et al., 2020).

Average accuracy and correct response times are plotted in **Figure 2A** and **2B**, respectively. According to a linear mixed effect model, the probability of a correct response varied significantly across presentation conditions (F(3, 10453)=6.26, p<0.001, BF=199) Planned contrasts revealed a significant drop in accuracy in the one-word trials compared to zero-word trials (F(1, 10453)=108.76, p<0.001, BF=3.09) and no significant difference between the one-word and two-word sequential trials (F(1, 10453)=2.95, p=0.08, BF=0.67). The former effect could be due to a bias to report “0” when uncertain. Importantly, there was a significant drop in the two-word simultaneous compared to two-word sequential trials (F(1,10453)=9.48, p=0.002, BF=6.53). That is a behavioral analogue of the simultaneous suppression effect. On two-word simultaneous trials in which participants made an error, they more often reported seeing one word (94% of total incorrect trials) than zero words (6% of incorrect trials).

**Figure 2:**
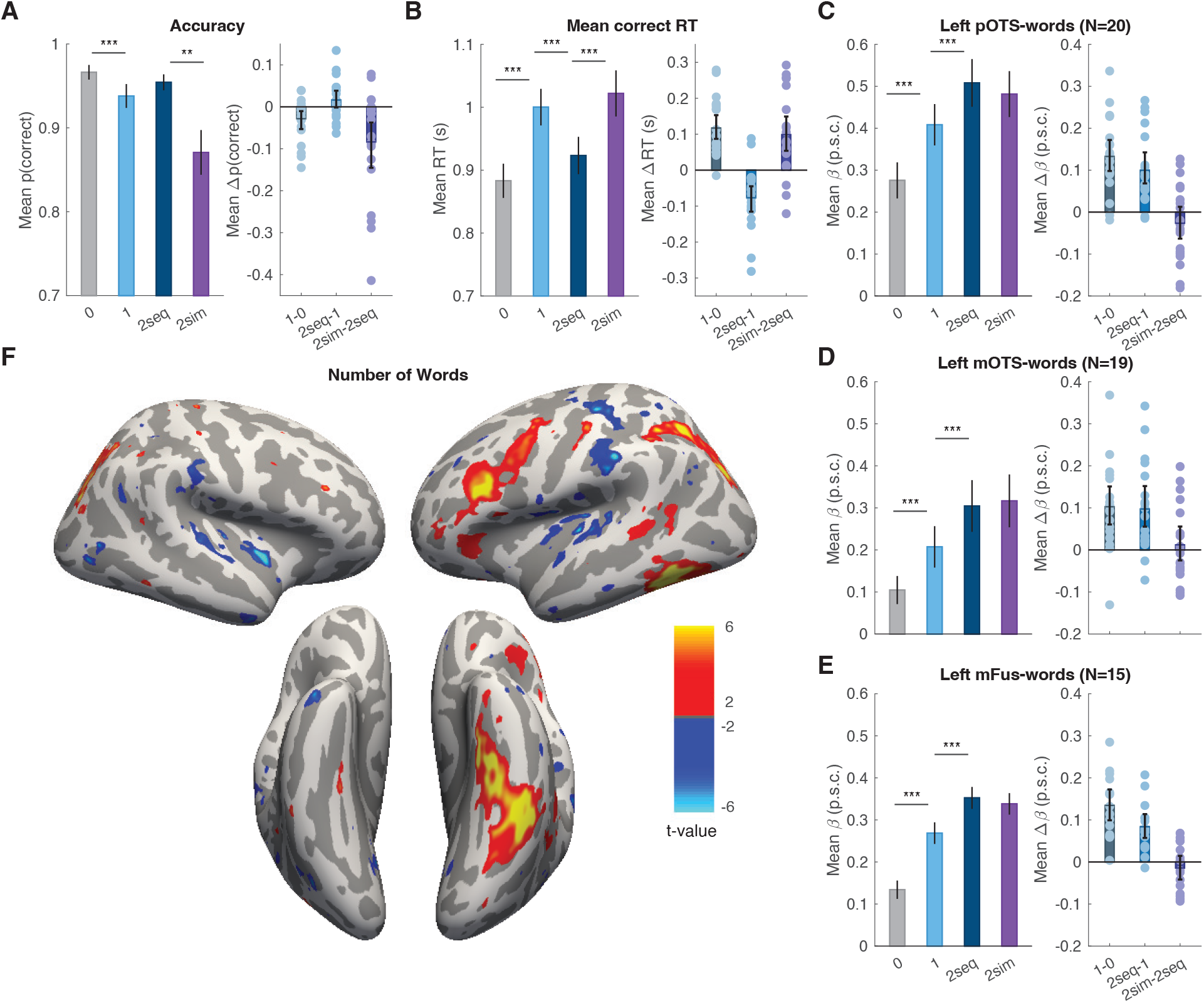
Effects of presentation condition on behavior and brain responses. **(A)** The left panel shows mean task accuracy for reporting the number of words as a function of the presentation condition. “2seq” means two words presented sequentially, while “2sim” means two words presented simultaneously. Error bars are `±` 1 SEM. Asterisks denote p-values from pairwise comparisons (***p<0.001, **=p<0.01, *=p<0.05). The right panel shows the *differences* in accuracy across pairs of conditions. The dots are individual participants, the bars are the mean differences, and the error bars are 95% bootstrapped confidence intervals. **(B)** Correct response times as a function of presentation condition, formatted in panel A. **(C, D and E)** Mean BOLD responses in three left ventral text-selection regions, plotted in units of percent signal change (p.s.c), and formatted as in panel A. **(F)** Whole-cortex maps of the effect of the number of words presented per trial. Colors represented t-values on the mean slope across participants, thresholded for p<0.05 with false discovery rate correction across all surface nodes. There are clusters of significant effect in the left inferior frontal sulcus, intraparietal sulcus, occipito-temporal sulcus and fusiform gyrus.

As explained in the Methods section above, we calculated response times (RTs) starting from the onset of frame 2, which for most conditions was the critical frame that revealed the correct answer. To ensure a fair comparison across conditions, we excluded from this analysis trials with two words in frame 1, or one-word trials in frame 1. There was a significant effect of presentation condition on correct RTs (F(3, 7339)=18.26, p<0.001, BF=2.34x10^5^). See **Figure 2B**. We observed significantly longer RTs for the one-word trials compared to zero-word trials (F(1, 7339)=482.64, p<0.001, BF=1.39x10^4^) and for one-word trials compared to two-word sequential trials (F(1, 7339)=17.75, p<0.001, BF=61.68). We also found a cost of simultaneous presentation: slower correct responses for two-word simultaneous compared to two-word sequential trials (F(1, 7339)=15.97, p<0.001, BF=45.90).

Taken together, the accuracy and RT results show that it was more difficult to detect two words when they were presented simultaneously than sequentially, which suggests a processing capacity limit (Scharff et al., 2011).

### Responses in ventral temporal text-selective areas increase linearly with the number of words but do not show simultaneous suppression

Our primary regions of interest lie in the occipito-temporal sulcus and the mid-fusiform gyrus of the left hemisphere. These text-selective regions have been shown to play a critical role in reading (Dehaene et al., 2015; Dehaene & Cohen, 2011). We expected these cortical regions to be among the few that could distinguish words from false font strings and would therefore show sensitivity to the number of words embedded in the sequences of false fonts.

fMRI and intracranial recordings have demonstrated that the VWFA is not unitary, but composed of subregions with different functions (e.g., Caffarra et al., 2021; Lerma-Usabiaga et al., 2018; White et al. 2019; Woolnough et al. 2021). We therefore defined three text-selective subregions from the same contrast in our visual category localizer: pOTS-words, mOTS-words, and mFus-words. A prior study implicated the left mOTS-words (also called VWFA-2) as a “bottleneck” that prevents simultaneous processing of two words (White et al., 2019). mFus-words has been less studied with fMRI, but has appeared in intracranial recordings as an important site anterior and medial of the “classical” VWFA (Woolnough et al., 2021). Within the ventral temporal cortex, this area is the first to respond differentially to low- vs high-frequency words. Woolnough and colleagues (2021) therefore propose that the mid-fusiform area supports the brain’s “orthographic lexicon” (see also Taylor et al., 2013).

The results for each of these three regions are reported separately in **Figure 2**, **Table 1** and **Table 2**. To maximize power, we begin with linear mixed-effect models that integrated single-trial data across these three regions and included region as a fixed effect.

**Table 1:** Statistical results from linear mixed effects models of two effects: the Number of Words effect and Simultaneous Suppression. The first modelled single-trial BOLD responses as a function of the number of words presented (excluding two-word simultaneous trials). The second tested for a difference in BOLD responses between two-word sequential and two-word simultaneous trials. We report the estimates of the slope for each fixed effect (in units of percent signal change), the standard error of that slope, its 95% bootstrapped confidence intervals, t-values, associated p-values corrected for multiple tests using FDR correction, and Bayes Factor for each effect. N is the number of subjects in which each ROI was present.

| Hemi | ROI | N | Experimental Condition | Slope | SEM | 95% CI | t value | p value | BF |
| --- | --- | --- | --- | --- | --- | --- | --- | --- | --- |
| LH | pOTS-words | 20 | Number of Words | 0.116 | 0.015 | [0.087, 0.145] | 7.88 | <0.001 | 1.46x10<sup>9</sup> |
|  |  |  | Simultaneous Suppression | 0.014 | 0.01 | [-0.006, 0.033] | 1.396 | 0.43 | 0.49 |
|  | mOTS-words | 19 | Number of Words | 0.1 | 0.016 | [0.069, 0.131] | 6.3 | <0.001 | 9.23x10<sup>5</sup> |
|  |  |  | Simultaneous Suppression | -0.005 | 0.01 | [-0.024, 0.015] | -0.482 | 0.63 | 0.28 |
|  | mFus-words | 15 | Number of Words | 0.109 | 0.011 | [0.088, 0.131] | 10.119 | <0.001 | 4.6x10<sup>9</sup> |
|  |  |  | Simultaneous Suppression | 0.008 | 0.007 | [-0.006, 0.022] | 1.065 | 0.43 | 0.39 |
| RH | pOTS-words | 11 | Number of Words | 0.008 | 0.005 | [-0.002, 0.017] | 1.62 | 0.16 | 0.15 |
|  |  |  | Simultaneous Suppression | 0.002 | 0.004 | [-0.007, 0.01] | 0.413 | 0.73 | 0.33 |
|  | mOTS-words | 7 | Number of Words | 0 | 0.01 | [-0.019, 0.019] | -0.004 | 1 | 0.08 |
|  |  |  | Simultaneous Suppression | 0.004 | 0.006 | [-0.007, 0.015] | 0.685 | 0.63 | 0.50 |

**Table 2:** Statistical results reporting pairwise differences for the number of words effect for left hemisphere text-selective regions. The third row for each ROI is a difference of differences in the first two rows. Here, we report the mean differences across conditions, the standard error, 95% bootstrapped confidence intervals, t-values, associated p-values corrected for multiple tests using FDR correction and Bayes Factor for each comparison.

| ROI | Comparison | Mean Difference | SEM | 95% CI | t value | p value | BF |
| --- | --- | --- | --- | --- | --- | --- | --- |
| pOTS-words | 1 - 0 | 0.133 | 0.019 | [0.098, 0.172] | 6.784 | <0.001 | 1.06x10<sup>4</sup> |
|  | 2seq - 1 | 0.1 | 0.018 | [0.068, 0.142] | 5.254 | <0.001 | 5.66x10<sup>2</sup> |
|  | 2seq - 1 vs. 1 - 0 | -0.033 | 0.023 | [-0.076, 0.018] | -1.374 | 0.19 | 0.53 |
| mOTS-words | 1 - 0 | 0.103 | 0.023 | [0.06, 0.151] | 4.371 | <0.001 | 88.5 |
|  | 2seq - 1 | 0.097 | 0.024 | [0.055, 0.152] | 3.937 | <0.001 | 38.1 |
|  | 2seq - 1 vs. 1 - 0 | -0.005 | 0.035 | [-0.061, 0.09] | -0.147 | 0.88 | 0.24 |
| mFus-words | 1 - 0 | 0.134 | 0.019 | [0.099, 0.172] | 6.964 | <0.001 | 3.19x10<sup>3</sup> |
|  | 2seq - 1 | 0.084 | 0.015 | [0.057, 0.114] | 5.504 | <0.001 | 3.58x10<sup>2</sup> |
|  | 2seq - 1 vs. 1 - 0 | -0.051 | 0.026 | [-0.101, 0.003] | -1.884 | 0.081 | 1.07 |

In these ventral temporal text-selective regions there was a significant positive effect of the number of words presented (0, 1 or 2-sequential) on the BOLD responses (F(1, 20346)=96.02, p<0.001, BF=5.27x10^9^), which did not interact with region (F(1, 20346)=0.21, p=0.81, BF=0.02). In this analysis, we also found a main effect of region (F(1, 20346)=5.83, p=0.003, BF=1.31x10^8^), as BOLD magnitudes were overall strongest in pOTS-words. The significant number of words effect was also present in each region analyzed separately (see Figure 2C-E and Table 1). The mean differences across each pair of conditions are plotted in the rightmost column of **Figure 2A-E**. The increase in responses from zero-word to one-word trials, one-word to two-word trials (all significant) are reported separately for each text-selective area in **Table 2**, as are the second-order comparisons of the 0-to-1-word increase versus the 1-to-2-word increase, none of which were significantly different. These results suggest that there is a steady increase in the response magnitude across trials with 0, 1 and 2 sequential words.

However, we did not find a simultaneous suppression effect in these text-selective regions. Integrating across all three regions, there was no difference between the 2-sequential and 2-stimultaneous conditions (F(1, 13559)=0.33, p=0.56, BF=0.13), and presentation condition did not interact with region (F(1, 13559)=2.09, p=0.12, BF=0.07). Statistics for the simultaneous suppression effect within each region are reported in Table 1, and individual subject differences shown as light purple dots in the right column of **Figure 2A-E**. Therefore, none of the text-selective regions in the ventral temporal cortex showed a simultaneous suppression effect that would mirror the effect observed in the behavioral data.

### Whole-cortex analyses reveal a language network that responds strongly to multiple words but no simultaneous suppression

A whole-cortex analysis revealed three large clusters in which the response increased significantly with the number of words presented. **Figure 2F** shows the map of t-values for the positive linear slope of the BOLD response with the number of words per trial, thresholded for FDR-corrected p<0.05. The first cluster was in the left ventral temporal cortex, covering parts of the occipito-temporal sulcus and the fusiform gyrus. This corresponds with the locations of the mOTS, pOTS and mFus in most subjects. The second cluster was in the left frontal cortex, including the inferior frontal sulcus (IFS) and extending posteriorly to the premotor cortex. This includes regions that are often referred to as “Broca’s area”, and it overlaps with some of the language ROIs that are analyzed separately below. The third cluster was in the intraparietal sulcus (IPS), bilaterally. These patches overlap with regions known to be involved in attentional control (Lauritzen et al., 2009; Silver et al., 2005). In the left hemisphere, this IPS region extended more anteriorly than in the right, including areas that may be more specifically involved in word recognition and language (Forseth et al., 2018; Ossmy et al., 2014; Rapp et al., 2016; Woolnough et al., 2022). All of these word-sensitive regions are known to show correlated activity during word recognition tasks (Chauhan et al., 2024; Stevens et al., 2017; Vogel et al., 2012; White et al., 2023).

We also performed a whole-cortex analysis of the simultaneous suppression effect: the difference in responses between two-word simultaneous and two-word sequential trials. After correction for multiple comparisons, we found no surface nodes with a significant effect.

### No simultaneous suppression in other visual or language areas

We also quantified the effects of the number of words and simultaneous suppression in seven other individually-defined regions, which we report in two sets: bilateral occipito-temporal visual regions, and the left frontal and temporal language regions. The first set includes: V4; the fusiform face areas (FFA-1 and FFA-2), and a limb-selective area (VTC-limbs). These results are plotted in **Figure 3**, and statistics are reported in **Table 3** and **Table 4**.

**Figure 3:**
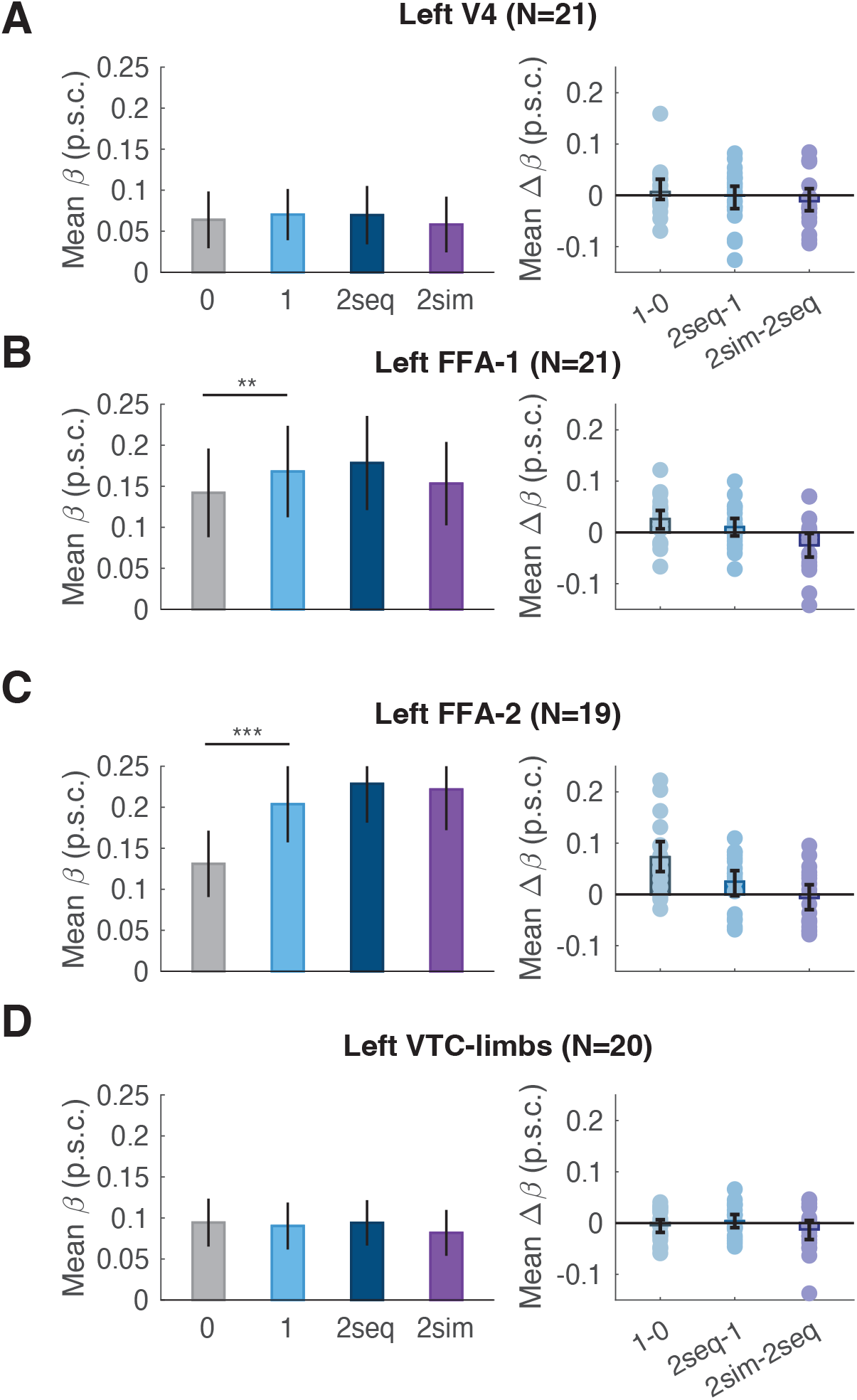
Effect of stimulus condition on BOLD responses in control visual ROIs. As in Figure 2, the left panels plot mean BOLD responses (with error bars that represent the SEM across subjects), and the right panels depict differences in responses between pairs of conditions (with error bars that represent bootstrapped 95% confidence intervals). The dots represent differences within individual subjects. The asterisks indicate p-values as Figure 2.

**Table 3:**
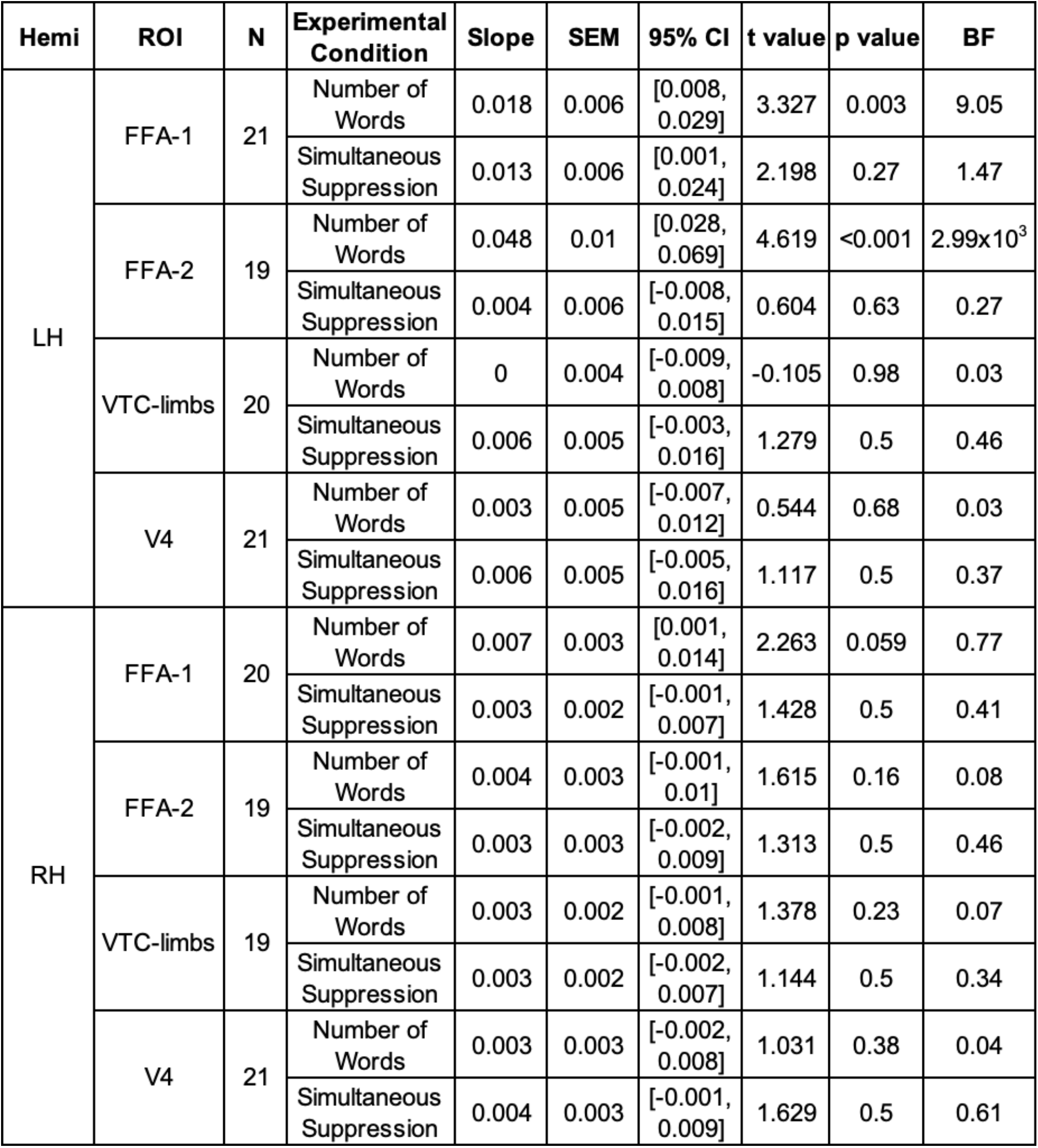
Statistics on control visual ROIs. Formatted the same as Table 1.

| Hemi | ROI | N | Experimental Condition | Slope | SEM | 95% CI | t value | p value | BF |
| --- | --- | --- | --- | --- | --- | --- | --- | --- | --- |
| LH | FFA-1 | 21 | Number of Words | 0.018 | 0.006 | [0.008, 0.029] | 3.327 | 0.003 | 9.05 |
|  |  |  | Simultaneous Suppression | 0.013 | 0.006 | [0.001, 0.024] | 2.198 | 0.27 | 1.47 |
|  | FFA-2 | 19 | Number of Words | 0.048 | 0.01 | [0.028, 0.069] | 4.619 | <0.001 | 2.99x10<sup>3</sup> |
|  |  |  | Simultaneous Suppression | 0.004 | 0.006 | [-0.008, 0.015] | 0.604 | 0.63 | 0.27 |
|  | VTC-limbs | 20 | Number of Words | 0 | 0.004 | [-0.009, 0.008] | -0.105 | 0.98 | 0.03 |
|  |  |  | Simultaneous Suppression | 0.006 | 0.005 | [-0.003, 0.016] | 1.279 | 0.5 | 0.46 |
|  | V4 | 21 | Number of Words | 0.003 | 0.005 | [-0.007, 0.012] | 0.544 | 0.68 | 0.03 |
|  |  |  | Simultaneous Suppression | 0.006 | 0.005 | [-0.005, 0.016] | 1.117 | 0.5 | 0.37 |
| RH | FFA-1 | 20 | Number of Words | 0.007 | 0.003 | [0.001, 0.014] | 2.263 | 0.059 | 0.77 |
|  |  |  | Simultaneous Suppression | 0.003 | 0.002 | [-0.001, 0.007] | 1.428 | 0.5 | 0.41 |
|  | FFA-2 | 19 | Number of Words | 0.004 | 0.003 | [-0.001, 0.01] | 1.615 | 0.16 | 0.08 |
|  |  |  | Simultaneous Suppression | 0.003 | 0.003 | [-0.002, 0.009] | 1.313 | 0.5 | 0.46 |
|  | VTC-limbs | 19 | Number of Words | 0.003 | 0.002 | [-0.001, 0.008] | 1.378 | 0.23 | 0.07 |
|  |  |  | Simultaneous Suppression | 0.003 | 0.002 | [-0.002, 0.007] | 1.144 | 0.5 | 0.34 |
|  | V4 | 21 | Number of Words | 0.003 | 0.003 | [-0.002, 0.008] | 1.031 | 0.38 | 0.04 |
|  |  |  | Simultaneous Suppression | 0.004 | 0.003 | [-0.001, 0.009] | 1.629 | 0.5 | 0.61 |

**Table 4:**
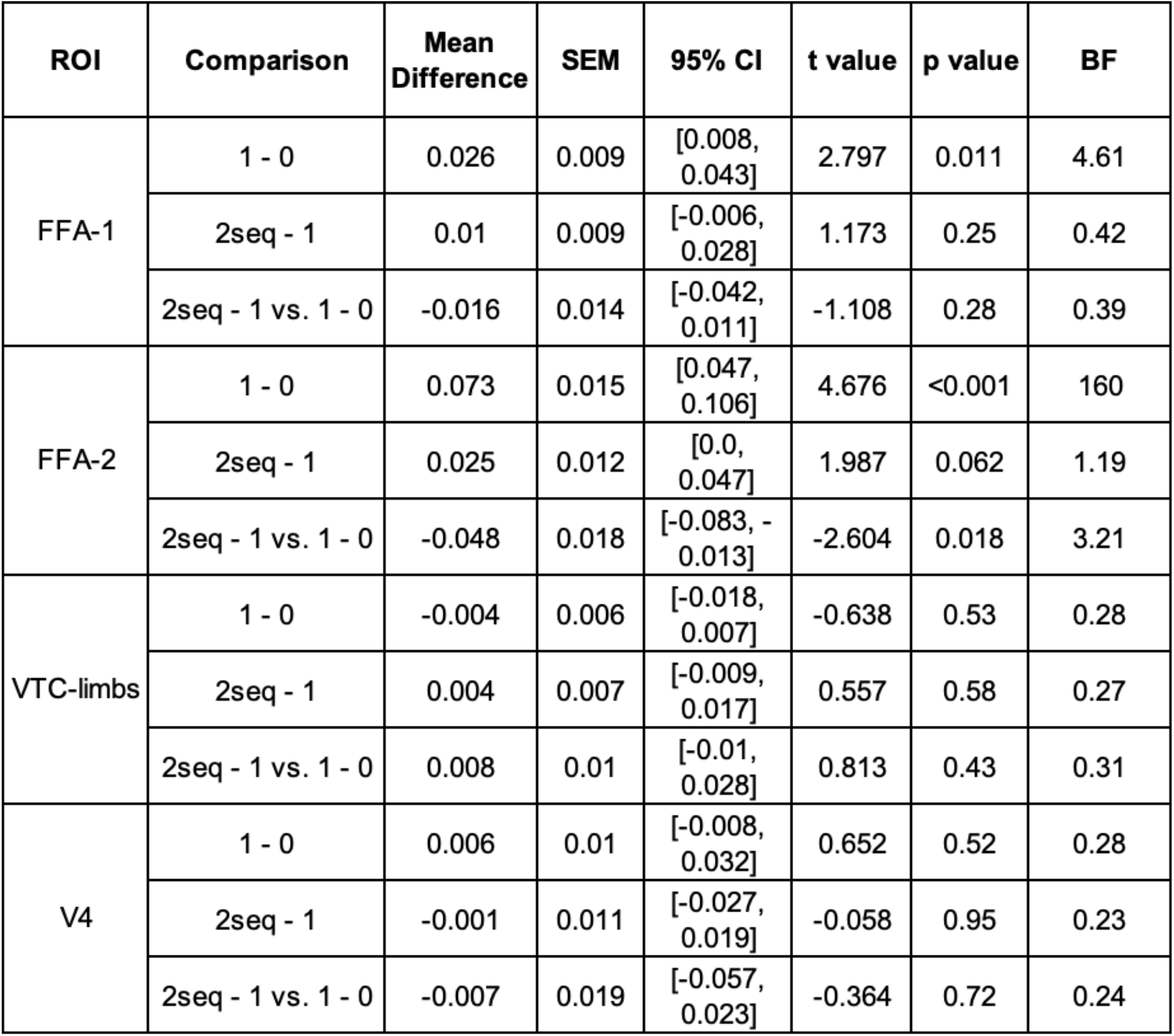
Pairwise differences between conditions in the left hemisphere control visual ROIs. Data formatted similarly to Table 2.

| ROI | Comparison | Mean Difference | SEM | 95% CI | t value | p value | BF |
| --- | --- | --- | --- | --- | --- | --- | --- |
| FFA-1 | 1 - 0 | 0.026 | 0.009 | [0.008, 0.043] | 2.797 | 0.011 | 4.61 |
|  | 2seq - 1 | 0.01 | 0.009 | [-0.006, 0.028] | 1.173 | 0.25 | 0.42 |
|  | 2seq - 1 vs. 1 - 0 | -0.016 | 0.014 | [-0.042, 0.011] | -1.108 | 0.28 | 0.39 |
| FFA-2 | 1 - 0 | 0.073 | 0.015 | [0.047, 0.106] | 4.676 | <0.001 | 160 |
|  | 2seq - 1 | 0.025 | 0.012 | [0.0, 0.047] | 1.987 | 0.062 | 1.19 |
|  | 2seq - 1 vs. 1 - 0 | -0.048 | 0.018 | [-0.083, -0.013] | -2.604 | 0.018 | 3.21 |
| VTC-limbs | 1 - 0 | -0.004 | 0.006 | [-0.018, 0.007] | -0.638 | 0.53 | 0.28 |
|  | 2seq - 1 | 0.004 | 0.007 | [-0.009, 0.017] | 0.557 | 0.58 | 0.27 |
|  | 2seq - 1 vs. 1 - 0 | 0.008 | 0.01 | [-0.01, 0.028] | 0.813 | 0.43 | 0.31 |
| V4 | 1 - 0 | 0.006 | 0.01 | [-0.008, 0.032] | 0.652 | 0.52 | 0.28 |
|  | 2seq - 1 | -0.001 | 0.011 | [-0.027, 0.019] | -0.058 | 0.95 | 0.23 |
|  | 2seq - 1 vs. 1 - 0 | -0.007 | 0.019 | [-0.057, 0.023] | -0.364 | 0.72 | 0.24 |

We focus first on the effect of the number of words in each trial. V4 and VTC-limbs were not sensitive to the number of words. Both left face-selective areas (FFA-1 and FFA-2) had responses that increased significantly with the number of words. The more anterior region, FFA-2, showed a strong significant increase in responses from the zero-word to one-word trials (BF=1.6e+02), but not the one-word to two-word sequential trials (Statistics on differences between conditions reported in **Table 4**). This partial sensitivity to the number of words in the left FFA-2, which we have observed before (Chauhan et al., 2024; White et al., 2023), could be due to overlap between primarily word-selective and primarily face-selective neural populations. Our ROI boundaries are somewhat arbitrary, it is unsurprising that the FFA shows a pattern of sensitivity to words that is similar to the neighboring VWFA, but weaker. None of the right hemisphere visual regions were affected by the number of words or simultaneous suppression (**Table 3**).

Importantly, none of these visual regions showed evidence of simultaneous suppression: responses were statistically indistinguishable between the 2-sequential and 2-simultaneous conditions, as in the text-selective regions (Statistics for all control visual ROIs reported in **Table 3**).

Results for the language regions are visualized in **Figure 4** and statistics are reported in **Tables 5 and 6**. These regions were defined in individual participants from a contrast of response to meaningful sentences versus meaningless pseudoword sequences (Fedorenko et al., 2010). First, we analyzed responses from three regions in the left frontal lobe. We found no evidence of simultaneous suppression in any of them, although they differed in how sensitive they were to the number of words presented on each trial. First, activity in the most dorsal region, the precentral gyrus (PG), did not vary significantly with the number of words. Second, a region in the left inferior frontal sulcus (IFS, **Figure 4B**) showed a significant number of words effect. Third, a region in the frontal operculum showed a very similar pattern of results to IFS.

**Figure 4:**
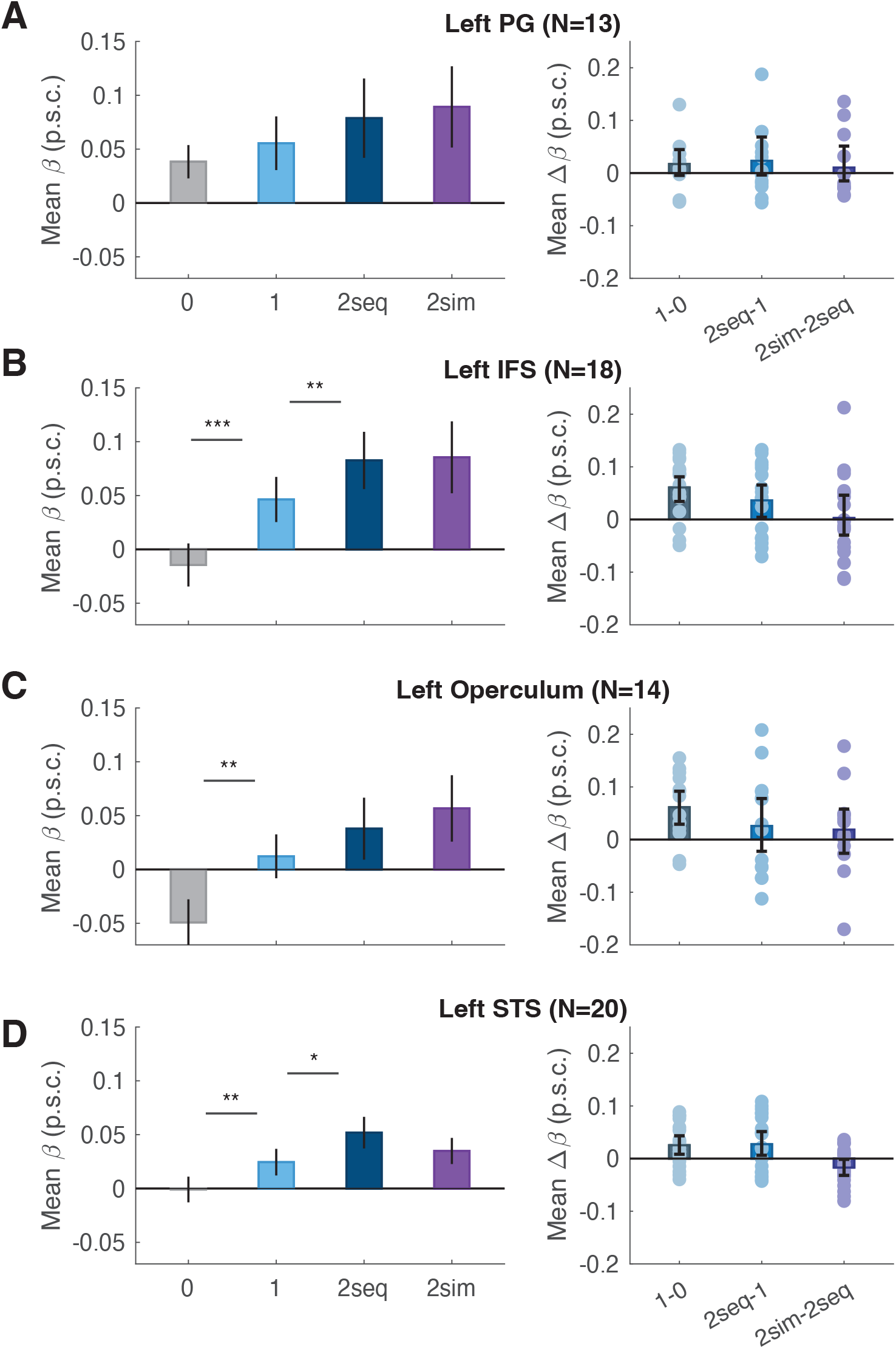
Effect of stimulus condition on BOLD responses in four language ROIs. Format as in Figure 3.

**Table 5:** Statistics on language ROIs. Formatted similar to Table 1.

| Hemi | ROI | N | Experimental Condition | Slope | SEM | 95% CI | t value | p value | BF |
| --- | --- | --- | --- | --- | --- | --- | --- | --- | --- |
| LH | PG | 13 | Number of Words | 0.02 | 0.012 | [-0.004, 0.044] | 1.671 | 0.16 | 0.38 |
|  |  |  | Simultaneous Suppression | -0.005 | 0.008 | [-0.02, 0.01] | -0.635 | 0.63 | 0.33 |
|  | IFS | 18 | Number of Words | 0.048 | 0.009 | [0.031, 0.066] | 5.442 | <0.001 | 1.6x10<sup>4</sup> |
|  |  |  | Simultaneous Suppression | -0.001 | 0.009 | [-0.019, 0.017] | -0.131 | 0.9 | 0.25 |
|  | Operculum | 14 | Number of Words | 0.043 | 0.014 | [0.016, 0.07] | 3.158 | 0.005 | 10.6 |
|  |  |  | Simultaneous Suppression | -0.01 | 0.011 | [-0.03, 0.011] | -0.934 | 0.58 | 0.37 |
|  | STS | 20 | Number of Words | 0.026 | 0.007 | [0.012, 0.041] | 3.564 | 0.002 | 85.2 |
|  |  |  | Simultaneous Suppression | 0.008 | 0.004 | [0.001, 0.016] | 2.103 | 0.27 | 1.75 |
| RH | STS | 17 | Number of Words | 0.006 | 0.003 | [0.0, 0.011] | 2.068 | 0.083 | 0.24 |
|  |  |  | Simultaneous Suppression | 0.002 | 0.002 | [-0.003, 0.006] | 0.643 | 0.63 | 0.28 |

**Table 6:** Statistics on pairwise condition in left hemisphere language ROIs. Formatted similar to Table 2.

| ROI | Comparison | Mean Difference | SEM | 95% CI | t value | p value | BF |
| --- | --- | --- | --- | --- | --- | --- | --- |
| PG | 1 - 0 | 0.017 | 0.01 | [-0.004, 0.048] | 1.34 | 0.21 | 0.58 |
|  | 2seq - 1 | 0.023 | 0.02 | [-0.004, 0.068] | 1.271 | 0.23 | 0.54 |
|  | 2seq - 1 vs. 1 - 0 | 0.006 | 0.02 | [-0.029, 0.048] | 0.305 | 0.77 | 0.29 |
| IFS | 1 - 0 | 0.061 | 0.01 | [0.033, 0.082] | 4.693 | <0.001 | 147 |
|  | 2seq - 1 | 0.036 | 0.02 | [0.006, 0.065] | 2.339 | 0.032 | 2.06 |
|  | 2seq - 1 vs. 1 - 0 | -0.025 | 0.02 | [-0.065, 0.02] | -1.109 | 0.28 | 0.41 |
| Operculum | 1 - 0 | 0.061 | 0.02 | [0.029, 0.091] | 3.629 | 0.003 | 15.2 |
|  | 2seq - 1 | 0.026 | 0.03 | [-0.021, 0.078] | 0.992 | 0.34 | 0.41 |
|  | 2seq - 1 vs. 1 - 0 | -0.036 | 0.03 | [-0.092, 0.033] | -1.072 | 0.3 | 0.44 |
| STS | 1 - 0 | 0.025 | 0.01 | [0.009, 0.042] | 2.926 | 0.009 | 5.77 |
|  | 2seq - 1 | 0.027 | 0.01 | [0.006, 0.05] | 2.321 | 0.032 | 2.00 |
|  | 2seq - 1 vs. 1 - 0 | 0.002 | 0.01 | [-0.024, 0.031] | 0.143 | 0.89 | 0.23 |

Interestingly, on zero-word trials (in which only false font strings were presented), the average response in the left frontal operculum dipped below the baseline BOLD level of 0 (by 0.05±0.02 p.s.c. on average; CI = [-0.009, 0.093], t(13)=2.21, p=0.045; BF=1.69). See the gray bar in **Figure 4C**. This is reminiscent of our previous finding that unfamiliar characters (false fonts) elicit suppression of the frontal language regions (but only when the participant is looking for familiar words; Chauhan et al., 2024).

Lastly, we identified a large language-sensitive region in the left superior temporal sulcus (STS; **Figure 4D**). The overall slope for the number of words effect was significant (**Table 5**). There was no effect on the number of words in the precentral gyrus. While the frontal operculum showed sensitivity to number of words, it was purely driven by the increase in response to a single word being present, compared to three frames of false fonts. Adding a second word did not significantly increase responses in these regions. IFS and STS showed a similar response as the ventral text-selective regions. Statistics to pairwise conditions illustrating this finding are reported in **Table 6**.

### Evidence of a capacity limit from lexical frequency effects

The fact that the mean BOLD responses were equivalent for simultaneous and sequential pairs of words could mean that reading-related brain regions can *detect* two strings of letters at once amidst the sequence of false font strings. However, that does not necessarily mean that two legible letter strings presented simultaneously were both processed *lexically* in the same way as two words presented in rapid succession were processed. The following exploratory analysis uses a signature of lexical processing to dig deeper into the comparison of the sequential and simultaneous trials, focusing on the ventral temporal text-selective areas.

Text-selective areas (often referred to as the VWFA) are known to respond more strongly to *less* common words (e.g., Chauhan et al., 2024; Kronbichler et al., 2004; Woolnough et al., 2021). There are a range of explanations for this effect, but it is generally agreed that low-frequency words require more time and/or effort for the word recognition system (Taylor et al. 2013; Gagl et al. 2022). If the VWFA acts as the “orthographic lexicon,” high-frequency words evoke lower responses there because of more rapid lexical access (Kronbichler et al., 2004; Wimmer & Ludersdorfer, 2018). Another explanation proposes that low-frequency words evoke larger VWFA responses because they are more difficult to distinguish from non-words, and again, that difficulty increases the total response (Gagl et al. 2022). Alternatively, low-frequency words elicit stronger “prediction errors” from higher-level language areas that feed back to the VWFA (Price & Devlin, 2011). All of these models attribute the lexical frequency effect to a component of the word recognition process that goes beyond simply detecting letters strings. We therefore analyze the magnitude of the lexical frequency effect to compare the degree to which such lexical processes were engaged in the simultaneous and sequential presentation conditions. This mirrors an approach taken by White et al. (2019), who reported that VWFA-2 (i.e., mOTS-words) was sensitive to the lexical frequency of only one attended word at a time.

We separately analyzed two sets of two-word trials: those in which both words were of low frequency (below the median of 4.1) and those in which both words were of high frequency (above the median). The results are plotted in **Figure 5**, for behavioral accuracy, response times and BOLD responses in left ventral text-selective areas. We compared the lexical frequency effects across the simultaneous and sequential presentation conditions.

**Figure 5:**
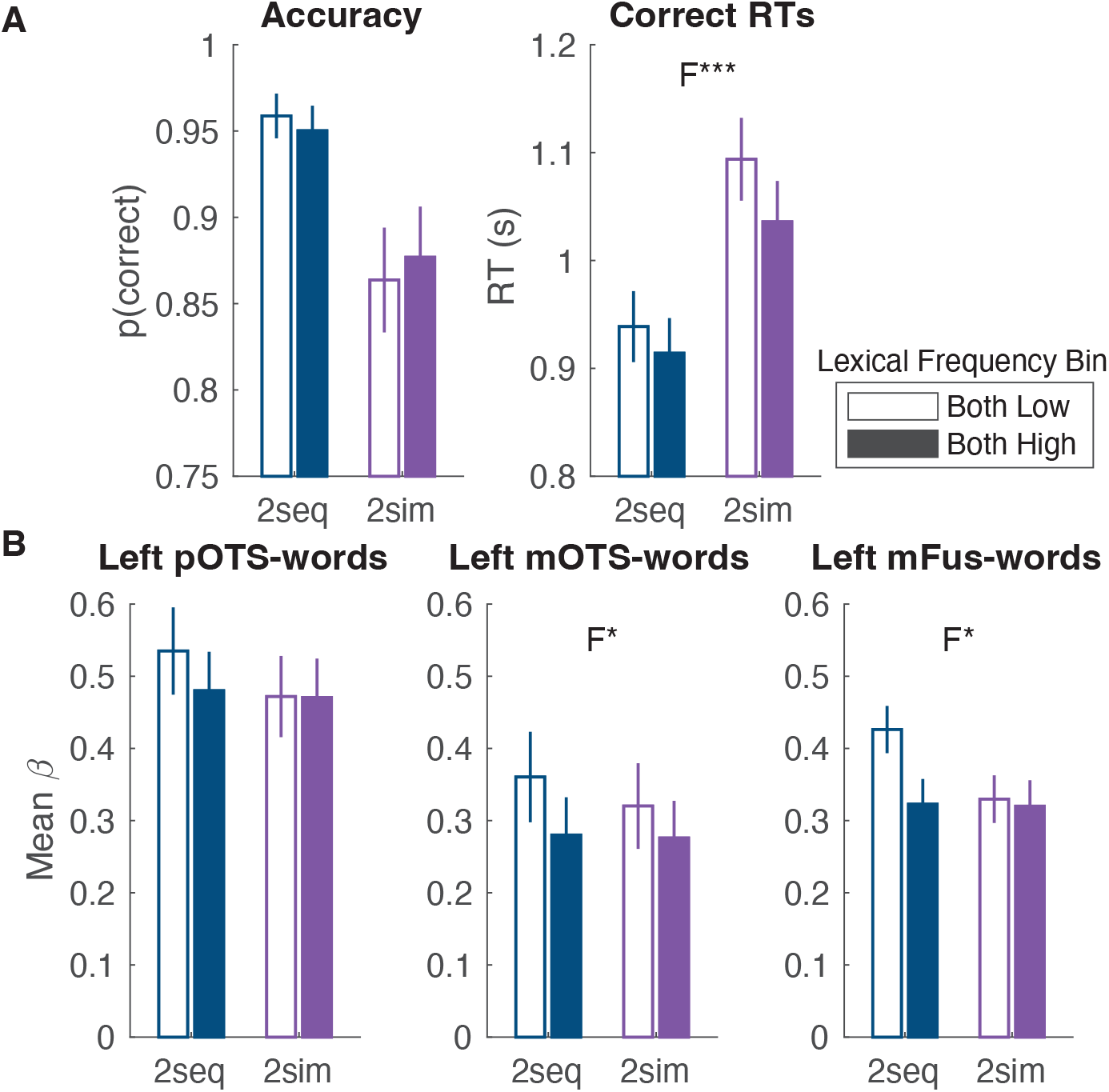
Lexical frequency effects on responses to two words in text-selective regions. **(A)** Behavioral results as a function of presentation condition and lexical frequency. “2-seq” means two words presented sequentially, “2-sim” means two words presented simultaneously. Filled bars are for trials with two high-frequency words, empty bars are for trials with two low-frequency words. **(B)** Average responses in text-selective ventral regions. Error bars are `±` 1 SEM. F indicates a significant main effect of lexical frequency. The asterisks indicate: ***=p<0.001, **=p<0.01, *=p<0.05.

Behaviorally, the probability of a correct response was lower on simultaneous than sequential trials (as reported in the main analysis above; F(1, 2641)=12.62, p<0.001, BF=407). Accuracy was not affected by lexical frequency bin (F(1, 2641)=0.25, p=0.62, BF=0.14), and the interaction between presentation condition and lexical frequency was not significant (F(1, 2641)=1.96, p=0.16, BF=0.22). Correct RTs were slower for simultaneous than sequential trials overall (F(1, 2409)=25.96, p<0.001, BF=3.8x10^7^), and slower for low-frequency than high-frequency word pairs (F(1, 2409)=11.59, p<0.001, BF=1.42), but there was no significant interaction between presentation condition and lexical frequency (F(1, 2409)=1.78, p=0.18, BF=0.29).

The BOLD responses in all three left ventral temporal text-selective areas are plotted in **Figure 5B**. To analyze these responses with maximal power, we fit a linear mixed-effect model that integrated across the three regions (with region as a fixed effect). The model outputs can be interpreted as a three-way ANOVA. We found no significant main effect of presentation condition (F(1,13547)=1.64, p=0.20, BF=0.45), but a main effect of lexical frequency bin (F(2,13547)=3.91, p=0.02, BF=1.54) and a main effect of region (F(2,13547)=5.13, p=0.006, BF=3.64x10^7^). The two-way interaction between region and lexical frequency bin was not significant (F(2, 13547)=0.84, p=0.49, BF=0.05).

Critically, the two-way interaction between presentation condition and lexical frequency bin was significant according to the *F* test (F(2, 13547)=5.20, p=0.006) although its Bayes Factor was low (BF=0.46).^1^ That interaction reflects this difference: there was a significant effect of lexical frequency bin in the sequential presentation condition (F(2, 6772)=7.22, p=7.40x10^-4^, BF=3.53), but not in the simultaneous condition (F(2,6775)=2.78, p=0.062, BF=0.161). That difference can be seen in **Figure 5B**: larger differences between responses to low- vs. high-frequency words in the sequential condition (“2seq”) than the simultaneous condition (“2sim”). The three-way interaction between region, lexical frequency bin and presentation condition was not significant (F(4, 13547)=0.80, p=0.52, BF=0.08).

Although the analysis above did not provide strong statistical evidence that these effects differed across the three regions, we also conducted exploratory analyses of the lexical frequency effect and its interaction with presentation condition in each region separately. We did so because the analysis above may be underpowered to detect a three-way interaction, and because prior studies suggest that lexical frequency effects are more indicative of a processing bottleneck in mOTS-words pOTS-words (White et al., 2019), and that mFus-words is the primary region where lexical frequency effects first arise (Woolnough et al, 2021). The p-values for the subsequent F tests (derived with linear mixed effect model fits) were FDR-corrected across the three regions.

In these separate analyses, the main effect of lexical frequency was not significant in pOTS-words (F(1, 2533)=2.47, p=0.12, BF=0.35), but was significant in mOTS-words (F(1, 2399)=5.75, p=0.025, BF=7.47) and mFus-words (F(1, 1935)=6.73, p=0.025, BF=2.45). The interaction between presentation condition and lexical frequency bin was not significant at the p<0.05 level in any of the individual regions, after correcting for the three comparisons (pOTS-words: F(1, 2533)=1.47, p=0.34, BF=0.43; mOTS-words: F(1, 2399)=0.39, p=0.53, BF=0.29; mFus-words: F(1, 1935)=4.07, p=0.13, BF=1.29). These results suggest that the effect of lexical frequency increases from pOTS-words to mOTS-words to mFus-words, which would be consistent with prior reports (White et al, 2019; Woolnough et al, 2021). Evidence for a reduction of the lexical frequency effect in the simultaneous presentation condition also appears strongest in mFus-words. However, we do not draw strong conclusions from these subsidiary tests because the three-way interaction between region, frequency bin and presentation condition was not significant, and even in mFus-words the statistical evidence for an interaction between frequency bin and condition was not strong.

Nonetheless, the analysis that integrated across all three regions found that overall, the relative elevation of BOLD responses to low-frequency words is weaker in the simultaneous condition than in the sequential condition. This result suggests that, despite the overall lack of simultaneous suppression shown in **Figure 2C-E**, simultaneous presentation did have a deleterious effect on lexical processing in the ventral temporal word-selective areas (**Figure 5**). For an explanation of this effect in terms of processing capacity limits, see the Discussion section.

### Serial and limited-capacity parallel models inspired by the lexical frequency effects

We now propose two simple models of how activity in the ventral text-selective regions could produce the pattern of lexical frequency effects. One model assumes rapid serial processing of words, and the other assumes limited-capacity parallel processing.

These models predict the magnitude of BOLD responses to pairs of words in the mFus-words region, where we find the strongest lexical frequency effects. See **Figure 6**. The core idea is that each word is processed until the accumulated “evidence” or “activation” reaches a threshold, or until the word(s) is replaced by another stimulus frame that interrupts processing. The total BOLD response is the integral of the activation over time for all words processed in that trial. This approach was inspired by Taylor, Rastle & Davis (2013). To account for the overall higher response to low- than high-frequency words, we assume that the evidence threshold is higher for low-frequency words. Thus, it takes more time to finish processing a low-frequency word, and the summed amount of activation is higher. This is roughly consistent with Bayesian accounts of word recognition (e.g., Norris, 2013) and previous proposals that low-frequency words are processed for more time or with more effort in the VWFA (Gagl et al. 2022; Kronbichler et al. 2004; Taylor et al. 2013). The free parameters in this model are: the slope of evidence accumulation over time, the low-frequency word threshold, and the high-frequency word threshold. The Python code that generates the model predictions is available along with our analysis code in the public repository for this study: https://doi.org/10.17605/OSF.IO/VXD9Z.

**Figure 6:**
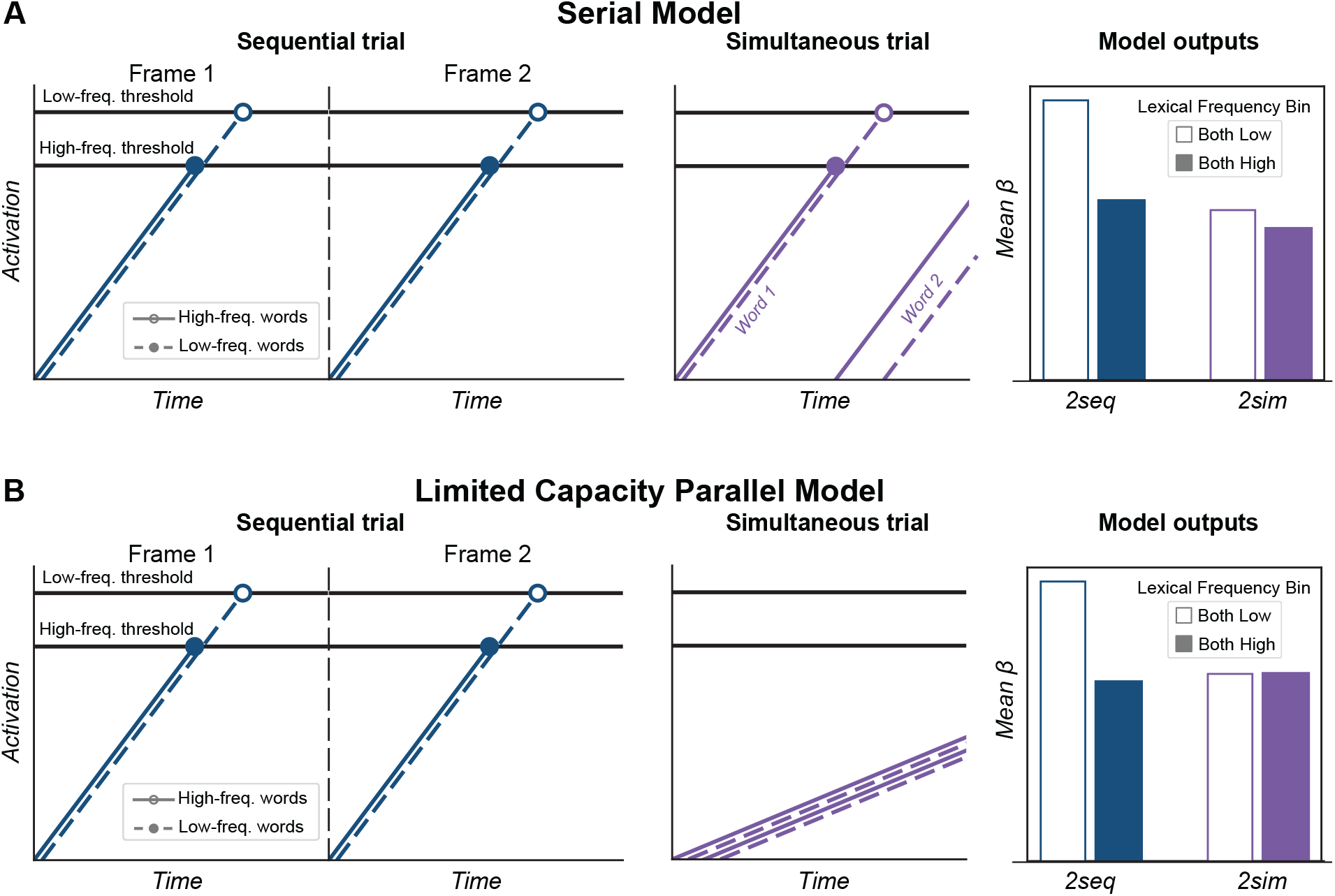
Two models that can explain the effects of the lexical frequency of word pairs in the sequential and simultaneous trials. Within each row, the first two panels show neuronal activation over time, as a form of evidence accumulation towards a word identification threshold. For both models, the threshold for low-frequency words is higher than for high-frequency words. We assume that on each trial either two low-frequency words are presented, or two high-frequency words are presented. The rightmost panels show the predicted BOLD responses, which are calculated as the integral of activation across time for each condition. Both models predict smaller effects of lexical frequency in the sequential condition than the simultaneous condition, as we observed empirically (Figure 5B). **(A)** Serial model: each word is processed one at a time. On simultaneous trials, the second word starts producing activation only after the first reaches its threshold, and the second word may not reach its threshold, which explains the behavioral deficit. **(B)** Limited-capacity parallel model: On simultaneous presentation trials, both words are processed simultaneously but with a shallower slope of activation over time than on sequential trials.

The serial model (**Figure 6A)** assumes that mFus-words processes one word at a time. If one word’s activation reaches its threshold before the next stimulus frame arrives, then processing of the second word begins immediately. In sequential trials (**Figure 6A**, upper two panels), one word is presented per frame and is fully processed (activation reaches the threshold). The area under the activation curve is larger for low- than high-frequency words (in **Figure 6A**, bottom right panel, compare the empty blue bar with the filled blue bar). In simultaneous trials (**Figure 6A**, bottom left panel), we assume that after processing one word, there is still time to switch to *begin* processing the other word, but only partially. When both words are low frequency, this switch occurs later and the second word does not get as close to its activation threshold, as compared to when both words are high-frequency. Simultaneous presentation reduces the relative boost in total response that occurs for low-frequency words, as compared to sequential trials when each word is fully processed (in **Figure 6A**, bottom right panel, compare the empty purple bar with the filled purple bar).

The limited-capacity parallel model (**Figure 6B**) assumes that mFus-words processes two words in parallel, but with a cost: two words presented simultaneously interfere with each other in the sense that both of their activations are *slowed*. The simulations for sequential trials are identical to the serial model, producing a clear lexical frequency effect (**Figure 6B**, top two panels). However, for the simultaneous trials, the slope of evidence accumulation for all words (i.e., activation over time) is shallower. That is the effect of the processing capacity limit, which could be accounted for by mutual inhibition among candidate lexical units that are activated by both words. Thus, neither of the words fully reach threshold before processing is interrupted, regardless of their frequency, and there is no difference between the total BOLD responses on high- and low-frequency trials (**Figure 6B**, bottom right panel, purple bars). Like the serial model, this model also correctly predicts that the responses to both high- and low-frequency words in the simultaneous condition are similar to the response to high-frequency words in the sequential condition.

Thus, it is possible to account for the lexical frequency effects with two different models. We did not quantitatively fit these simple models to the data, but we offer each as a proof of concept for a mechanistic account of activity in text-selective regions. Importantly, both models assume that that there is some kind of processing capacity limit. One assumes that there is a serial bottleneck in lexical identification, but that two high-frequency words can be at least partially processed in sequence within a single frame. The other assumes that there is parallel activation for two words at once, but with interference that manifests as a significant slowing of evidence accumulation. Figure 6 shows that both models predict similar patterns as we observed in the data, but they are not identical. The parallel model, given its assumption that the interference is so great that no words reach their activation thresholds in the simultaneous condition, predicts 0 lexical frequency effect in that condition. The serial model, given the parameter settings that allow one word to reach its activation threshold on each trial and another to begin processing, will always predict a small lexical frequency effect in the simultaneous condition, because of the higher threshold for low-frequency words that is reached for at least one word. However, our data do not clearly distinguish between the serial model’s prediction of a small effect and the parallel model’s prediction of 0 effect in the simultaneous condition. Thus, we are not attempting to draw any conclusions from the minor differences in the model outputs for the two models.

Future studies could explore stimulus and task conditions for which these two models make clearly different predictions. For example, if the duration of each stimulus was shorter, with subsequent masks that interrupt processing appearing sooner, then the model predictions diverge. The limited-capacity parallel model would predict a more rapid decrease in total response in the simultaneous condition. Other approaches could manipulate spatial attention to one or both words to distinguish parallel from serial processing (similar to White et al., 2019).

For now, we can reject an unlimited-capacity parallel model, which would assume that two words presented simultaneously are processed in the same way (with the same slope, reaching the same thresholds) as when two words are presented sequentially. Such a model would predict equivalent BOLD responses and lexical frequency effects in both presentation conditions.

## DISCUSSION

We first established that in ventral temporal text-selective regions, and in frontal and temporal language-processing regions, BOLD response magnitudes steadily increased with the number of words presented, from 0 to 1 to 2. Behaviorally, participants were less likely to detect both of two words that were presented simultaneously than sequentially, suggesting a processing capacity limit (Scharf et al., 2011; White et al., 2020; White et al., 2026). The neural correlate of that capacity limit could have presented as a weaker BOLD response to two words presented simultaneously as compared to sequentially, as occurs for other types of stimuli in visual cortex (Kaiser et al., 2014; Kastner, Weerd, et al., 2001; Miller et al., 1993; Shim et al., 2013). However, we did not observe statistically significant simultaneous suppression in any of our regions of interest nor in a whole-cortex analysis. These data demonstrate that text-selective regions can respond to two written words simultaneously.

However, there is a distinction between simply *detecting* two letter strings and identifying them as specific words in the mental lexicon. We also found that in ventral temporal text-selective areas, the response was higher for two low-frequency words than two high-frequency words, but less so in the simultaneous condition. The statistical evidence in this analysis was modest and must be followed up with further research. Our preliminary interpretation is that lexical access was impaired by the simultaneous presentation, resulting in less sensitivity to the words’ frequencies. We proposed two simple models to account for those effects, either assuming serial processing or parallel processing with interference (**Figure 6**).

We now consider four factors of our empirical design that may explain the absence of a neural simultaneous suppression effect:

1. *Visual Stimulation:* We equated the numbers of characters and stimulus onsets and offsets across conditions. Had we not done so, but simply presented a variable number of words without the ‘filler’ false font strings, we may have found a simultaneous suppression effect. However, such an effect could be due to nonlinear temporal integration of responses to stimulus transients, rather than competition for limited resources (Kupers et al. 2024).
2. *Stimulus Duration:* Each word was presented for 183 ms, which is short enough to minimize goal-directed eye movements within a frame. The interstimulus interval was 233 ms, roughly the average fixation duration during reading (Rayner et al., 2016). This interval is also not far from the 250 ms used in classic simultaneous suppression studies (e.g., Kastner et al., 1998; Kastner & Ungerleider, 2001; McMains & Kastner, 2011). However, there is some evidence that the effect increases as stimulus durations increase (Kupers et al., 2024). We chose not to make the time between frames any longer because that could invite eye movements. But we cannot definitively rule out shifts of *covert* attention between words within a single frame (see **Figure 6A**). If subjects attempted that strategy, however, they were not entirely successful, because task accuracy was impaired on simultaneous trials. Nonetheless, we might have found stronger simultaneous suppression had the presentation durations been even *shorter,* to prevent (partial) sequential processing of simultaneously presented words.
3. *Stimulus Location:* Unlike many past studies (e.g., Kastner et al. 1999; Kupers et al. 2024), we presented just two stimuli, each 1.5° from fixation. Both were inside the large population receptive fields of voxels in the ventral temporal cortex that cover roughly the central 5° of the visual field (Le et al., 2017). Therefore, we expected a strong effect in ventral temporal cortex (Kastner, De Weerd, et al., 2001; Kupers et al., 2024). To minimize the distance between words, we presented them above and below fixation. If anything, we would be *less* likely to find simultaneous suppression with the words presented to the left and right of fixation, which would be more familiar to English readers.
4. *Top-down attention:* While early studies found the strongest simultaneous suppression effect when the stimuli are ignored (Kastner et al., 1998; S. McMains & Kastner, 2011), it is also present when the stimuli are attended (Scalf et al., 2011). As our question is whether two words can be *read* simultaneously, we made our stimuli task-relevant. Voluntary effort to read amplifies activity in the VWFA, possibly via top-down signals from language regions (Chauhan et al., 2024; Kay & Yeatman, 2017; Mano et al., 2013; White et al., 2023). Thus, it is possible top-down signals counteracted a local simultaneous suppression effect that we might have observed if the stimuli were ignored. Conversely, an effect might have emerged had the task required further semantic processing. This concern is mitigated by two findings: there was a behavioral deficit in the simultaneous condition, and higher-level language regions were engaged and sensitive to the number of words.

### Comparison to a prior report of a serial bottleneck in the VWFA

White et al. (2019) measured BOLD responses to two words while participants performed a semantic categorization task and either focused attention on one word or divided attention between both. They concluded that the “serial bottleneck” apparent in task performance arises in mOTS-words (i.e., VWFA-2).

On the one hand, these findings led us to predict a simultaneous suppression effect in text-selective areas, which we did not find. Note that White et al. (2019) displayed words to the left and right of fixation and found a bias: the word on the right was easier to recognize and dominated VWFA activity. Our display arrangement avoided that complication, perhaps revealing more balanced processing of two words at once. On the other hand, our analyses of lexical frequency are consistent with White et al. (2019) in suggesting that the VWFA cannot process two words at once as well as it can process one word at a time.

### A hybrid parallel-then-serial model to account for behavioral and neural effects

To account for all the fMRI and behavioral data in this study, and to be consistent with several previous lines of research, we propose a hybrid model: sublexical information from two words is processed in parallel in word-selective cortex, followed by a serial process of identifying each word as a lexical item (**Figure 7**). The parallel stage is consistent with an existing model of reading that proposes that multiple words are processed in parallel at the stage of activating letter or bigram detectors (Grainger et al., 2014, 2016; Snell et al., 2018; Snell & Grainger, 2019). Orthographic information from different spatial locations is pooled into a single “channel” that activates multiple candidate word representations, which inhibit each other.

**Figure 7:**
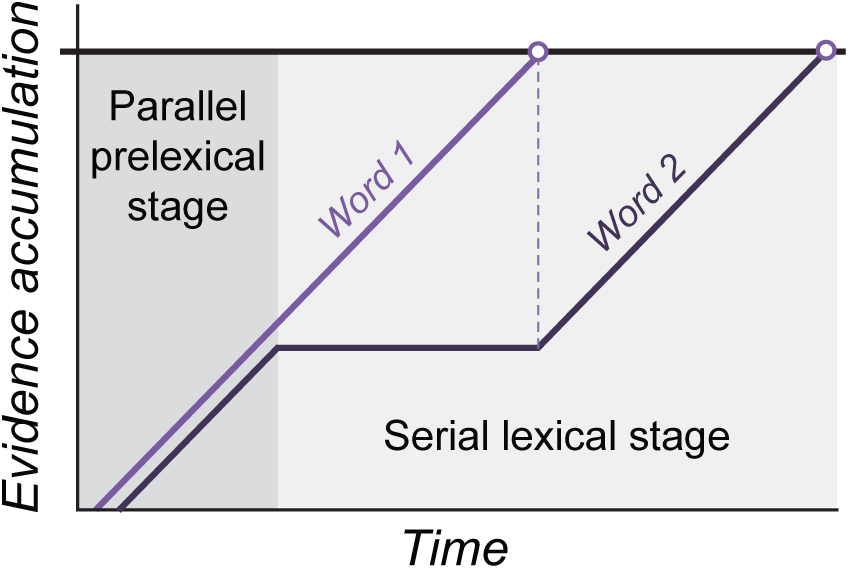
Schematic of a hybrid parallel-then-serial model for the processing of two simultaneously presented words. The logic of this graph is similar to the models in Figure 6 that are explained in the Results section, except that it does not assume processing is interrupted by a post-mask; hence this figure shows both words reaching the threshold for lexical access.

Building on those ideas, Anupindi et al. (2026) proposed a hybrid parallel-serial interactive activation model to account for behavioral data in tasks with attention divided between two words. They found large deficits for recognizing two *unrelated* words at once (one above and one below fixation), consistent with the serial model. But when two words formed a compound word, performance exceeded the serial model. This suggests some degree of parallel processing that can maintain information about one word while another is identified.

We propose that when two words are presented simultaneously, ventral-temporal text-selective areas encode sublexical orthographic information about both words in parallel. However, further lexical processing occurs serially, perhaps relying on communication with higher-level language regions. Therefore, the sublexical evidence for one word is maintained while the other word rises to the activation threshold, and then, if there is sufficient time, lexical processing begins for the second word (hence the jagged shape of the blue line in **Figure 7**). Evidence accumulation ends when subsequent stimuli arrive that interrupt processing. The fact that processing of one word often does not complete causes the task performance deficits on simultaneous trials. However, both words contribute equal amounts of activation (the areas under the red and blue curves are equal), meaning that the total BOLD response is equivalent to when two words are presented sequentially.

### Future directions

An important question for future research is about when BOLD responses in the text-selective regions asymptote as the number of words increase. And is that threshold the same for all regions? The hybrid model we present raises the question of which brain regions participate in the transition from parallel sublexical processing to the selection of one word to be identified serially. It will be important to build upon this study with tasks that are more similar to natural reading in a variety of languages. For instance, the capacity for parallel processing might vary as a function of where the words are positioned, that that might differ across readers of scripts that are arranged vertically or horizontally. Building towards natural reading, our results also present avenues for exploring how responses to multiple words depend on their linguistic relationships: such as pairs of words that are semantically related or form a compound or a syntactic phrase.

## ACKNOWLEDGEMENTS

This study was funded in part by the National Eye Institute (grant R00 EY-029366). We thank Kimya Firoozan and Yen-Chu Lin for assistance with data collection.

## DATA AVAILABILITY

All raw and preprocessed data, as well as code for stimulus presentation and data analysis, are available on OpenNeuro.org (https://openneuro.org/datasets/ds007442) and OSF (https://doi.org/10.17605/OSF.IO/VXD9Z).

## Footnotes

^1^This is the only statistical test in which we found that the null-hypothesis test (an F test derived from a linear mixed effect model) led to a different conclusion than the Bayes Factor. It could be because unlike the F test, the Bayes Factor calculation was conducted on data first averaged over trials for each participant and did not allow for random slopes across participants.

